# An end-to-end in silico framework for truth-based validation and regression testing of clinical genome sequencing pipelines, from simulated reads to reportable findings

**DOI:** 10.64898/2026.11.01.763475

**Authors:** Bohuslav Dvorský, Mariana Komárková, Kateřina Kollinová, Ondřej Klempíř, Radim Krupička

## Abstract

Clinical genome sequencing pipelines change often, and each change requires proportionate revalidation. Guidelines accept in silico data as a supplement but leave variant interpretation outside pipeline validation, and, to our knowledge, no framework tests a pipeline against a known truth up to the reportable finding, so report specificity is not measured. We present a source-available framework that tests a pipeline end to end, from sequencing reads to the report, against a truth fixed before any run. It plants expert-curated pathogenic and benign variants from the same genes, and optionally synthetic variants that no database holds, into the GIAB HG002 germline background and records the expected class and report decision of each variant. It then simulates reads and runs secondary and tertiary analysis as black boxes through exchangeable adapters. Findings are compared with the truth at the call-set and report levels, each failure is assigned to the first pipeline stage where it appears, and a variant-by-variant comparison of two runs serves as a regression test. We applied the framework to nf-core/sarek with DeepVariant and to the ACMG/AMP classifier BIAS-2015, using 243 ClinGen-curated variants in five test sets. Single-nucleotide variant recall against the HG002 benchmark was 99.5% on simulated and 99.3% on real reads, and every planted variant was present in the reads and called with the correct genotype. The report included 115 of 117 pathogenic and likely pathogenic variants (98.3%) but withheld only 106 of 126 variants that must not be reported (84.1%), mainly because variants of uncertain significance were upgraded and recessive carriers were reported. Without ClinVar-derived criteria, report sensitivity fell to 74.4%. Most false reports of one test set came from diploid calling of the male chromosome X, and haploid calling removed them without side effects. In supplementary experiments, indels were called completely up to 20 bp, and BIAS-2015 left 74% of synthetic loss-of-function variants unreported because it combined criteria by the 2015 rules. The framework measures report-level sensitivity and specificity and localizes regressions, so it can supplement validation on human samples and support change control.

## 1 Introduction

Clinical genome sequencing relies on complex bioinformatic pipelines that transform raw sequencing reads into clinically reportable findings. The reliability of the final result depends on the performance of the individual tools used and how their outputs are combined across successive stages. These stages include read alignment, variant calling, filtering, annotation, and classification according to the recommendations of the American College of Medical Genetics and Genomics and the Association for Molecular Pathology (ACMG/AMP; Richards et al., 2015) or classifications in curated databases. The final stage is the selection of findings for the clinical report (Rehder et al., 2021). Errors can propagate across stages and alter the final interpretation (Lavrichenko et al., 2025). The Sequencing Quality Control 2 study found that the bioinformatic workflow had the largest impact on interlaboratory reproducibility of germline variant detection (Mercer et al., 2021). Furthermore, as software, parameters, and reference databases evolve, pipelines require appropriate verification or revalidation (Roy et al., 2018). However, revalidation practices vary substantially. In a 2019 AMP survey of 61 participants, 16% reported revalidating more than twice a year, while 15% reported never revalidating (Duncavage et al., 2023). These challenges underscore the need for a systematic, reproducible framework that tests clinical sequencing pipelines against a predefined standard of truth, from raw reads to the final reporting decision. Such a framework should also support regression testing after updates and identify the stages responsible for discrepancies. Here we present such a framework and apply it to an open-source clinical pipeline, from simulated reads to the report.

Validation guidelines cover secondary analysis in more detail than the interpretation and reporting of variants. The recommendations of the Association for Molecular Pathology and the College of American Pathologists (AMP/CAP) end with a list of variant calls carrying basic annotation (Roy et al., 2018). The Nordic Alliance for Clinical Genomics left annotation outside its scope (Lavrichenko et al., 2025), and the joint recommendations on in silico data leave detailed recommendations for tertiary analysis outside theirs (Duncavage et al., 2023). Both AMP/CAP and the in silico recommendations nevertheless expect the later stages to be tested. Automated annotation must be thoroughly validated (Roy et al., 2018), and an interpretation pipeline should not be validated on known pathogenic ClinVar variants alone (Duncavage et al., 2023). In silico data have a defined but limited role in this testing. They may not replace end-to-end validation with human samples (Roy et al., 2018, Recommendation 14; Duncavage et al., 2023, Recommendation 1), but they may be used to test minor pipeline updates (Hume et al., 2019; Rehder et al., 2021; Duncavage et al., 2023, Recommendation 4). In practice, update testing is still limited. All Nordic units test new releases on previously analyzed samples, an approach biased toward recall at the expense of precision, and only one unit runs automated tests on every commit, merge, and release. The units also recognize an over-reliance on Genome in a Bottle (GIAB) truth sets, which are often used both to train and to validate tools (Lavrichenko et al., 2025).

Existing in silico approaches test either the sequencing data or the interpretation, but not both against one truth. De novo simulators such as VarSim (Mu et al., 2015) and NEAT (Stephens et al., 2016) generate reads from a diploid genome carrying the chosen variants, so the whole pipeline runs from raw reads. Purely simulated data, however, are not recommended for the critical phase of assay validation, because they may overestimate performance (Duncavage et al., 2023). Read-editing tools such as BAMSurgeon (Ewing et al., 2015), insiM (Patil et al., 2019), and VarBen (Li et al., 2021) instead insert variants into reads from a real laboratory run, which keeps the error profile of the assay (Duncavage et al., 2016). insiM, for example, selects uniquely mapped reads at each target locus, mutates enough of them to reach the requested allele fraction, and writes new FASTQ files together with a Variant Call Format (VCF) file of the introduced variants (Patil et al., 2019). Because editing relies on correct prior mapping, it usually cannot assess highly homologous genes such as *PMS2* and *SMN1*/*SMN2* (Duncavage et al., 2023), and it does not test the aligner unless edited files are converted back to FASTQ (Furtado, 2024). Edits can also fail: in one germline study, seven of 49 selected variants could not be used because insiM placed them at the wrong position (Roy et al., 2025). Evaluations of both families typically stop at variant-level outputs, although interpretation is itself a substantial source of uncertainty. Among 2,419 variants classified by at least two of 12 laboratories, 34.4% were discordant on the five-tier scale but only 3.2% on a two-tier actionable scale (Mighton et al., 2022). Against expert classifications from the Clinical Genome Resource (ClinGen), the best of four automated tools matched exactly for 58.4% of 256 variants (Costa et al., 2025). Such benchmarks start from curated variant lists, so they do not capture errors that arise before classification (Costa et al., 2025; Eisenhart et al., 2025). Bastida-Lertxundi et al. (2025) came closer to the full chain by uploading FASTQ files with 24 known causal variants to seven commercial platforms, where strict concordance with the reference class ranged from 42% to 75%. All 24 variants were pathogenic or likely pathogenic (P/LP), however, and the authors call for studies that also include population variants. The real-world methodology of Mitev (2026) assigns each observation to the layer where it arose, but its cohorts were enriched for pathogenic variants, so it could not estimate specificity. To our knowledge, no framework tests a pipeline against a predefined truth from raw reads to the reporting decision, measures report-level specificity on known benign controls, and attributes each discrepancy to the stage where it arose. Such attribution matters because a single control spanning several steps leaves open where an error occurred (Jennings et al., 2017).

To address these gaps, we have developed a tool-agnostic in silico framework for truth-based validation and regression testing of clinical genome sequencing pipelines. Its unit of testing is the expected finding: a planted variant together with its expected ACMG/AMP class or laboratory curated class, the classes accepted as correct, and the expected report decision. The framework makes three contributions. First (C1), it validates a pipeline end to end, from FASTQ files to the report decision. Benign or likely benign (B/LB) variants planted in the same genes as the P/LP variants serve as negative controls, so report-level specificity can be measured on this defined set. This differs from variant-calling specificity, which standard benchmarking omits because true negatives are hard to define (Krusche et al., 2019). Second (C2), it extends end-to-end validation to change control. Revalidation after pipeline changes is required (Roy et al., 2018), and regression testing has been recommended (Lavrichenko et al., 2025); here it runs against predefined expected outcomes across secondary and tertiary analysis, and each discrepancy between runs is attributed to the stage where it arose. Third (C3), it uses de novo diploid-genome simulation (Mu et al., 2015; Stephens et al., 2016) rather than read editing. This lets us confirm that every planted variant is present, tests the aligner, allows highly homologous regions, and needs no patient data; the cost is limited realism, which we quantify against real data. Components are exchanged through adapters, so laboratories can test their own tools. Because test sets are defined in configuration, laboratories can also build their own sets and repeat the same tests under different conditions. We position the framework as supplemental validation and change control in the sense of Roy et al. (2018, Recommendations 14 and 17) and Duncavage et al. (2023, Recommendation 1); it does not replace validation with human samples.

## 2 Methods

### 2.1 Design principles and framework overview

The framework tests a clinical sequencing pipeline against a predefined truth: the expected variants, classes, and report decisions are fixed before any reads are simulated (Figure 1). The workflow consists of six steps: (1) selection of test variants; (2) encoding of the truth in a phased VCF file with expected-outcome metadata; (3) planting of the variants into a background genome and simulation of paired-end reads; (4) alignment and variant calling by the secondary pipeline; (5) annotation, classification, and reporting by the tertiary tool; and (6) comparison of the outputs with the truth at the VCF and report levels, with each discrepancy assigned to the stage where it arose.

**Figure 1.**
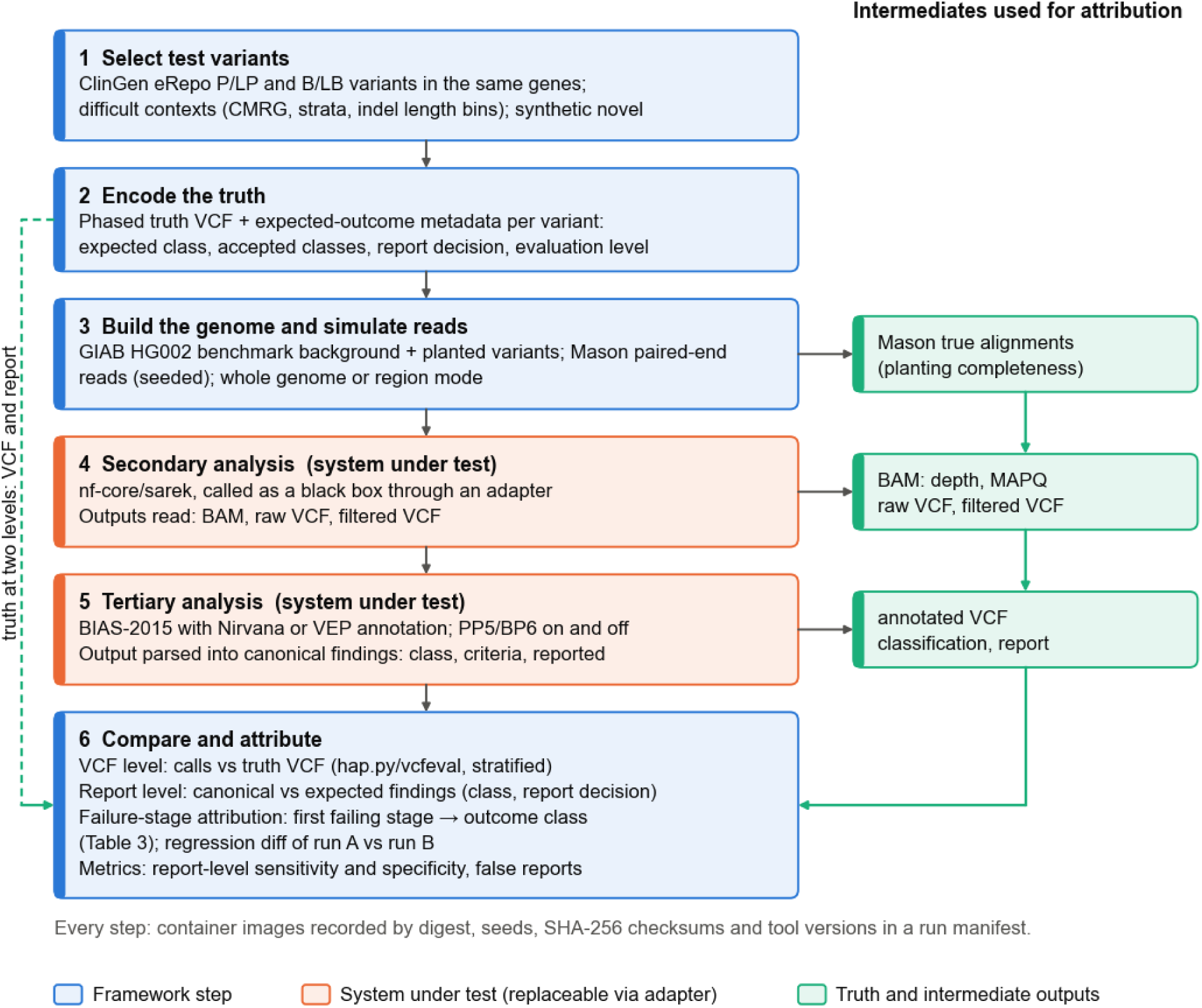
Overview of the framework. Steps 1–6 run in order. Steps 4 and 5 are the systems under test, reached only through adapters; alternatives to each component are listed in **Table 1**. The truth encoded in step 2 is compared with the outputs at two levels, VCF and report. The intermediate outputs on the right are used to assign each failure to the stage where it arose (**Table 3**; Figure 3A).

**Table 1.**
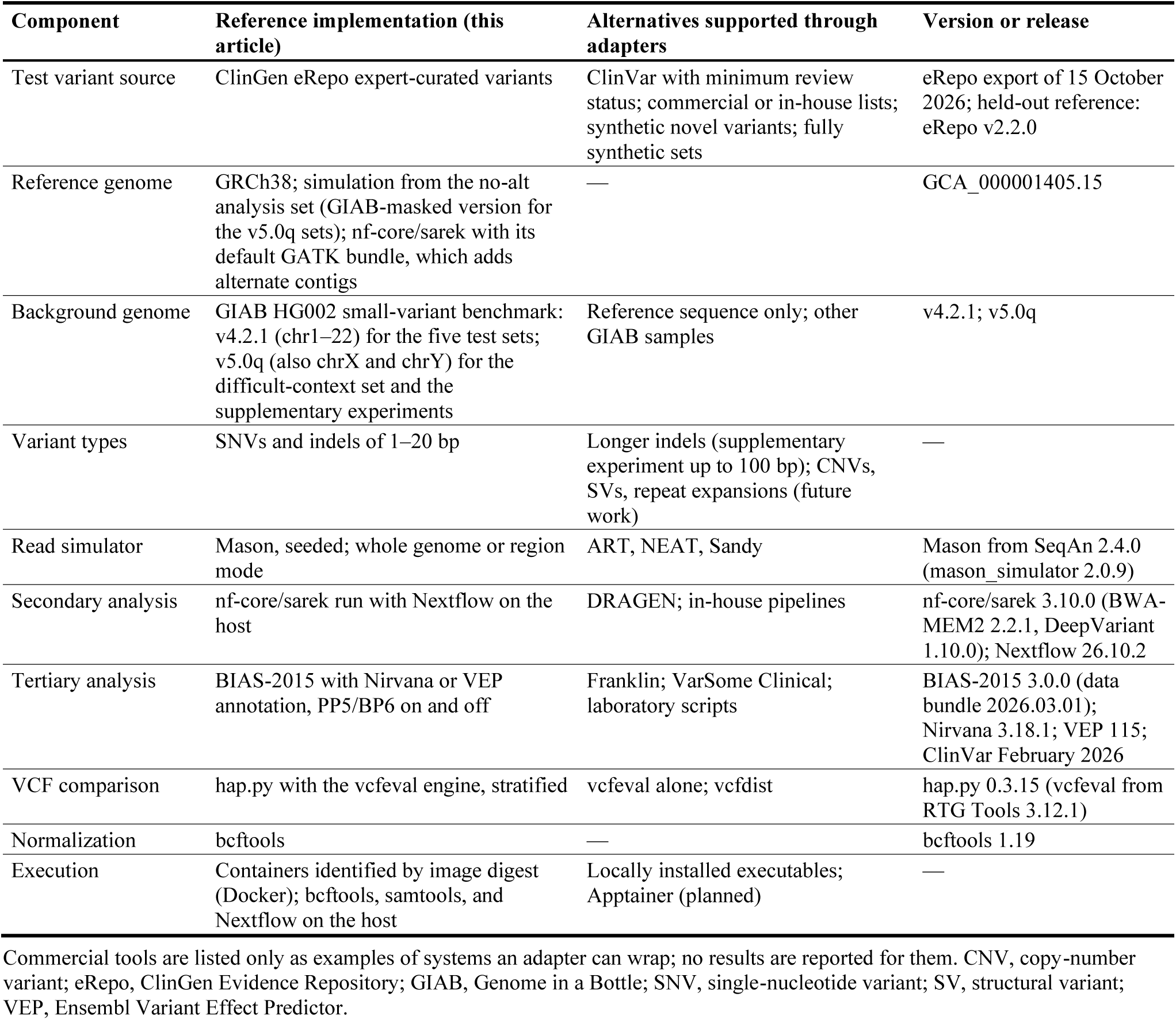
Framework components: reference implementation and supported alternatives.

The design follows four principles. First, the truth is known by construction, so no result depends on an orthogonal assay or on agreement between callers. Second, every planted variant is scored as an expected finding, which ties it to its expected class and report decision, so calling and interpretation are judged on the same variants. Third, each system under test is reached only through a declared interface, so a laboratory can replace one component without touching the others (Table 1). Fourth, all random choices derive from a seed and all tools run from container images recorded by digest, so the framework adds no run-to-run variation of its own. Variation from the systems under test is measured separately (see Metrics and acceptance criteria).

### 2.2 Selection of test variants

Scoring interpretation requires an expected class for every planted variant, assigned by experts rather than derived from any tool under test. Curated test variants therefore come from the ClinGen Evidence Repository (eRepo), which holds the classifications approved by ClinGen variant curation expert panels (VCEPs) together with the criteria they applied (Preston et al., 2022). The ClinGen variant curation process is recognized by the U.S. Food and Drug Administration, and classifications approved by its VCEPs are considered valid scientific evidence that can streamline test development and validation (Preston et al., 2022). We use eRepo rather than ClinVar (Landrum et al., 2018) because ClinVar aggregates submissions of variable quality without validation (Costa et al., 2025), whereas eRepo also records the evidence behind each classification (Preston et al., 2022). Records are mapped to GRCh38 from their genomic Human Genome Variation Society (HGVS) expressions and left-normalized. Alleles whose records disagree across classification tiers are excluded, and planted variants are kept a minimum distance apart. Table 2 lists the eRepo release, the genes, and the number of variants per category. Curated P/LP variants form the first category.

**Table 2.** Categories of test variants: sources, selection rules, numbers in this study, and expected outcomes.

| **Category** | **Source (release)** | **Selection** | **Variant types** | **n** | **Expected outcome** |
|---|---|---|---|---|---|
| Curated P/LP (positive) | ClinGen eRepo, export of 15 October 2026; held out = absent from the eRepo v2.2.0 benchmark of BIAS-2015 (lifted to GRCh38) | Rules by inheritance (autosomal dominant, X-linked, autosomal recessive biallelic); seeded ranking within genes; 27 genes | SNVs, indels 1–20 bp (planted: 1–5 bp) | 117 (P 51, LP 66) | Class P or LP (both accepted); reported; level: both |
| Curated P/LP, recessive carriers | ClinGen eRepo, export of 15 October 2026 | One P/LP variant in an autosomal recessive gene (GAA, RPE65), negative sample | SNVs, indels 1–20 bp (planted: 1–5 bp) | 6 (P 4, LP 2) | Class P or LP; not reported (one allele in a recessive gene); level: both |
| Curated B/LB (negative controls) | ClinGen eRepo, export of 15 October 2026; held out as above | Same genes as P/LP, same VCEPs where possible; 19 genes | SNVs, indels 1–20 bp (planted: 1–5 bp) | 86 (B 48, LB 38); held out 19 | Class B or LB (both accepted); not reported; level: both |
| Curated VUS (negative controls) | ClinGen eRepo, export of 15 October 2026; held out as above | Same genes as P/LP; 15 genes | SNVs, indels 1–20 bp (planted: 1–5 bp) | 34; held out 10 | Class VUS; not reported; level: both |
| Difficult contexts (framework option) | Variants in GIAB CMRG regions and GIAB stratifications (homopolymers, tandem repeats, segmental duplications) | Per stratum; indel ladder 1–5, 6–10, 11–20 bp | SNVs, indels by ladder bin | 292 in a separate test set (difficult_v5q); 450 in a supplementary experiment (Note S2) | Curated variants: as above; ClinVar-derived: no expected class, level: calling |
| Synthetic novel (framework option) | Explicit list in the suite configuration; absent from ClinVar and eRepo | Nonsense, canonical splice-site and frameshift variants in LoF genes; last exon avoided | SNVs, small indels | None in the five test sets; 324 in a supplementary experiment (Note S1) | Class P or LP, fixed before the run from a written rationale; reported; level: both |
| Background (not planted) | GIAB HG002 small-variant benchmark, GRCh38: v4.2.1, chr1–22 (five test sets); v5.0q, chr1–22, X and Y, PASS records of up to 20 bp in the benchmark regions (difficult-context set, supplementary experiments) | All records except those within the minimum distance of a planted variant (removed and logged) | SNVs, indels | v4.2.1: about 4.1 million records per whole-genome sample; 33–74 removed per sample | No class; level: calling; any reported background variant is a false report |

n, number of planted variants over the five test sets of this study (short1, 33; short2, 30; short3, 23; full1, 88; full2, 69); each curated category targets at least 59 variants, by analogy with the AMP/CAP minimum per variant type (Roy et al., 2018). VUS (34), held-out P/LP variants (36) and single P/LP variants in recessive genes (6) fall below this target, so their estimates have wider intervals. Held out: 36 of the 123 P/LP variants of both P/LP rows (6 further P/LP variants of unknown status), 19 B/LB and 10 VUS variants. B, benign; CMRG, Challenging Medically Relevant Genes; eRepo, ClinGen Evidence Repository; GIAB, Genome in a Bottle; LB, likely benign; LoF, loss of function; LP, likely pathogenic; P, pathogenic; SNV, single-nucleotide variant; VCEP, variant curation expert panel; VUS, variant of uncertain significance. Evaluation level: calling, interpretation or both (see Truth encoding).

**Table 3.** Outcome classes and failure-stage definitions.

| No. | Outcome class | Meaning | Stage and evidence |
|---|---|---|---|
| 1 | not_simulated | Planted variant absent from the simulated reads; a framework error, not counted against the system under test | Simulation: Mason true alignments |
| 2 | not_covered | Insufficient simulated depth at the locus | Simulation: depth of true alignments at the locus |
| 3 | mismapped | Allele not called and reads carrying it not aligned to the true locus, e.g. to a paralog | Alignment: BAM compared with true alignments |
| 4 | not_called | Variant absent from the raw VCF | Variant calling: raw VCF |
| 5 | filtered | Variant in the raw VCF but removed or with FILTER other than PASS | Filtering: filtered VCF |
| 6 | genotype_mismatch | Variant called with the wrong zygosity | Calling: genotype compared with the truth VCF |
| 7 | not_annotated | Variant lost or wrongly annotated during annotation | Annotation: annotated VCF |
| 8 | misclassified | Assigned class outside the accepted classes | Classification: canonical finding |
| 9 | report_mismatch | Class accepted, but the report decision differs from the expected one | Reporting: canonical finding (reported yes/no) |
| 10 | correct | All scored layers correct | — |
| — | false_report | Reported finding that matches no planted variant, e.g. an HG002 background variant or an off-target finding; counted separately from specificity | Reporting: report |

Classes are assigned in pipeline order, and the first failing stage determines the class. Only the layers given by the evaluation level of an expected finding are scored. A planted B/LB variant that is reported fails at classification (misclassified) or at reporting (report\_mismatch). Each outcome record also keeps the evidence at the locus (for example true-read depth, mapped depth, MAPQ, presence in raw and filtered VCF). Class names are those used in the code.

Curated variants can overlap the knowledge base on which a tertiary tool was tuned. In that case, the tool could reproduce the truth without applying the classification criteria. BIAS-2015 tuned its evidence thresholds on eRepo v2.2.0, which is also its benchmark (Eisenhart et al., 2025). Variants absent from that release are therefore flagged as held out and reported separately. The selection rules also accept other sources, such as ClinVar with a minimum review status, in-house lists, and synthetic variants.

Flagging held-out variants requires the list of variants in eRepo v2.2.0. The export itself is not published, but its variants are: the benchmark file of BIAS-2015 (Additional file 2 of Eisenhart et al., 2025) holds 8,707 records on hg19 from the snapshot accessed on 2 February 2025; the article itself reports 8,703 variants. We lifted the file to GRCh38 with CrossMap and the UCSC hg19-to-hg38 chain (Zhao et al., 2014); none of the 8,707 records failed. A planted single-nucleotide variant (SNV) is held out when its left-normalized allele is absent from the lifted file. Indels are matched by position instead: an indel counts as present when the file has any indel starting within 3 bp of its span. Position-based matching is used because many indel alleles in the file do not follow standard VCF representation; for example, multi-base deletions appear as single-base deletions without an anchor base. The rule is therefore conservative: it can only reduce, not inflate, the number of indels flagged as held out.

Because the variant list is reconstructed, we checked the held-out status against the approval date of the eRepo records. Of the 243 planted variants, 71 were absent from the benchmark file. Six of them, all P/LP, had been approved before the snapshot; we report them as of unknown status. The remaining 65 are held out: 36 of 123 P/LP, 19 of 86 B/LB, and 10 of 34 variants of uncertain significance (VUS; Table 2). Conversely, 29 variants present in the file had been approved again after the snapshot, which is expected for curations that a panel revises.

The second category consists of curated B/LB variants. They are selected from the same genes as the P/LP variants (Figure 2A) and, where possible, from the same VCEPs, for example *BRCA1* and *BRCA2*. Sensitivity and specificity are then measured on the same transcripts, annotation sources, and gene-level rules. Because reporting policies exclude B/LB variants from clinical reports (Miller et al., 2021; Durkie et al., 2024), they serve as negative controls for the report. Including both classes also meets the requirement that a validation set represent the expected clinical significance of variants rather than benign polymorphisms alone (Roy et al., 2018).

**Figure 2.**
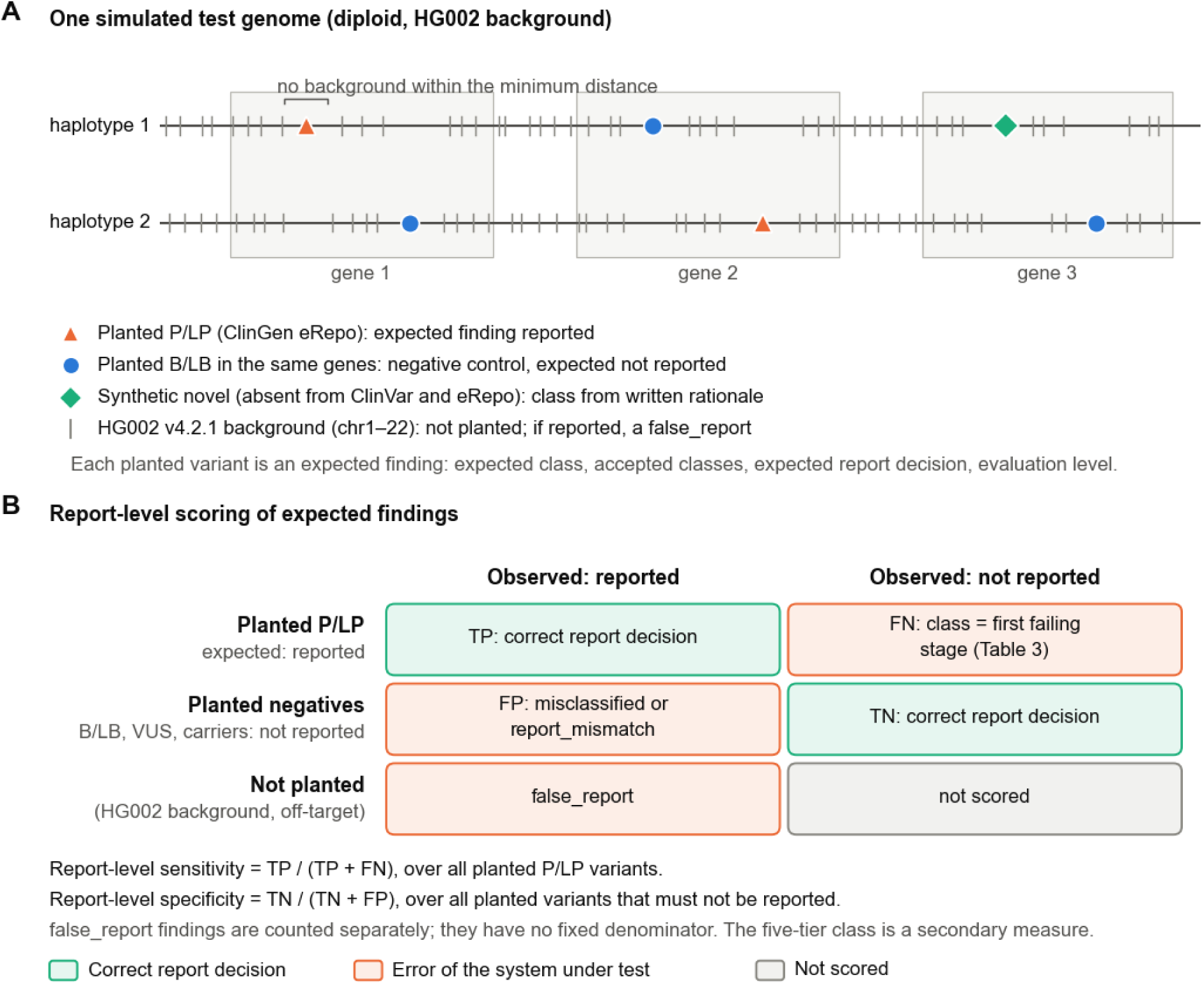
Test genome design and report-level scoring. (A) Planted variants in one simulated diploid genome (schematic, not to scale). P/LP and B/LB variants are planted in the same genes. Synthetic novel variants, a framework option used only in a supplementary experiment (Supplementary Note S1), require the classification criteria to be applied. The background consists of the germline SNVs and indels of the GIAB HG002 v4.2.1 benchmark (chromosomes 1–22); records within 500 bp of a planted variant are removed, and chromosome X carries no background. (B) Expected versus observed report decision. Planted P/LP variants give report-level sensitivity and planted variants that must not be reported (B/LB variants, VUS, and single P/LP variants in recessive genes) give report-level specificity. Reported findings that match no planted variant are counted separately as false reports. A failed expected finding carries the outcome class of its first failing stage (Table 3).

A third category targets contexts in which short-read calling is known to fail. Variants are drawn from the GIAB Challenging Medically Relevant Genes (CMRG) benchmark, which characterizes 273 of 395 autosomal genes that earlier benchmarks excluded because of repeats or polymorphic complexity (Wagner et al., 2022b). Further variants come from genome stratifications for homopolymers, tandem repeats, and segmental duplications (Dwarshuis et al., 2024). Such a dedicated set makes pipeline testing more realistic than the standard GIAB truth set alone (Lavrichenko et al., 2025). Rules can also select indels in length bins, for example 1–5, 6–10, and 11–20 bp. Indels longer than 15 bp remain challenging to call in medically relevant genes (Wagner et al., 2022b), and the largest indel a pipeline detects has to be established experimentally (Hume et al., 2019). A separate test set of this study used this category without length bins, and a supplementary experiment used length bins up to 100 bp (see Test sets used in this study).

The framework also accepts a fourth category: synthetic nonsense, canonical splice-site, and frameshift variants absent from ClinVar and eRepo, in genes where loss of function is an established disease mechanism. No database holds a classification for them, so the tertiary tool has to apply the ACMG/AMP criteria, principally PVS1 and PM2, rather than retrieve an assertion. The effect of database lookup can be large: switching off the ClinVar-based PP5 criterion in an earlier version of BIAS-2015 reduced its P/LP sensitivity on eRepo from 92.26% to 81.80% (Eisenhart et al., 2024). Positions where PVS1 needs caution, such as the last exon (Durkie et al., 2024), are avoided. A generator script draws such variants from a seed. It places them on MANE Select transcripts (Matched Annotation from NCBI and EMBL-EBI; Morales et al., 2022) of genes with established haploinsufficiency, keeps positions where the change is predicted to trigger nonsense-mediated decay, excludes positions recorded in ClinVar, eRepo, or gnomAD (v4.1 exomes, released 19 April 2024; Chen et al., 2024), and confirms the consequence with the Ensembl Variant Effect Predictor (VEP; McLaren et al., 2016; Supplementary Note S1). Each synthetic variant carries a written rationale and an expected class fixed before any run. We expected P or LP. The ClinGen Sequence Variant Interpretation (SVI) Working Group, which lowered PM2 to supporting strength, classifies PVS1 with PM2 at supporting strength as likely pathogenic (ClinGen Sequence Variant Interpretation Working Group, 2020), and the point-based guidelines of the Association for Clinical Genomic Science (ACGS), which keep PM2 at moderate strength, give such a variant 10 points, which is pathogenic (Durkie et al., 2024). The primary score is the report decision; a secondary analysis accepts a VUS based on PVS1 and PM2 alone, to separate errors in the criteria from errors in combining them.

The reference implementation is limited to SNVs and indels of 1–20 bp, within the scope of the AMP/CAP pipeline guideline (SNVs, indels, and delins of 21 bp or less; Roy et al., 2018). Copy-number and structural variants and repeat expansions are left to future work. AMP/CAP requires at least 59 variants of each type, such as SNVs and indels, to demonstrate 95% reliability at 95% confidence (Roy et al., 2018); we use the same number as the target for each category of test variants. Smaller categories are flagged and reported with correspondingly wider confidence intervals.

### 2.3 Test sets used in this study

We built five test sets from genes of the ACMG secondary-findings list, version 3.2 (Miller et al., 2023a; Table 2). The selection rules include all 29 of the 81 genes on the list that had at least one eRepo curation in the export used. Only curations of conditions for which the list recommends reporting were eligible; for RYR1, for example, malignant hyperthermia was eligible but RYR1-related myopathy was not. All planted variants of these sets are curated eRepo variants; the difficult-context and synthetic categories and the indel length bins described above were not used in them. Each set has two male samples. The positive sample carries P/LP variants that must be reported. The negative sample carries B/LB variants, VUS, and single P/LP variants in genes with autosomal recessive inheritance. None of these may be reported: the secondary-findings policy excludes VUS (Miller et al., 2021) and requires two P/LP variants in a recessive gene (Miller et al., 2023a). Every gene with a planted B/LB variant also carries planted P/LP variants (19 of 27 genes).

The five sets differ in size so that they serve different uses. Three short sets cover selected chromosomes (chr17 and chr19; chr12–chr15; chr1, chr10, and chrX) and suit frequent regression runs. Two whole-genome sets differ in how P/LP variants are drawn: a seeded random draw (full1) or one variant per gene (full2).

A sixth test set (difficult_v5q) covers difficult contexts. It holds 292 variants in two male samples: 267 ClinVar variants with at least two review stars, 60 each in segmental duplications, low-mappability regions, homopolymers, and tandem repeats, 20 in regions that GRCh38 falsely duplicates, and 7 in paralogous genes (SMN1, PMS2, and OTOA), scored at the calling level only; and 25 curated eRepo variants (15 P/LP and 10 B/LB) in difficult regions of secondary-findings genes, scored at both levels. Its background is the newer HG002 v5.0q benchmark, derived from the complete diploid HG002 assembly (Hansen et al., 2026), which also covers chromosomes X and Y, and its reads are simulated from a GRCh38 sequence in which GIAB masks the falsely duplicated copies (Behera et al., 2023), so reads come only from the true copy. The pipeline keeps the standard reference. Because the background differs, its results are reported separately.

Two supplementary experiments tested the remaining options of the selection rules in the setting of the difficult-context set. The indel-length experiment planted 450 ClinVar indels with at least two review stars in six length bins (1–5, 6–10, 11–20, 21–35, 36–50, and 51–100 bp; 66–80 per bin) in two male samples. Of these, 85 P/LP and 35 B/LB indels in secondary-findings genes were scored at both levels and the others at the calling level only. Indels longer than 20 bp lie outside the scope of the reference implementation; they show where calling stops. The synthetic experiment planted 324 generated variants in 33 secondary-findings genes in four male samples: 30 nonsense, 30 canonical splice-site, and 264 frameshift variants of 1–20 bp (169 insertions and 95 deletions). Supplementary Notes S1 and S2 describe both experiments; their results are reported separately.

### 2.4 Truth encoding: expected-outcome metadata

For each simulated sample, the framework writes a single-sample truth VCF on GRCh38, normalized with bcftools (Danecek et al., 2021). If normalization would change a planted record, the run stops. Genotypes are phased and assigned per sample: compound heterozygotes are placed in cis or in trans, and variants on chromosome X or Y outside the pseudoautosomal regions are hemizygous in male samples. The file carries only the GT and PS fields, so it contains no read-level values that no simulation has yet produced. INFO fields record the provenance of each variant: source database and release, record identifier, VCEP, and held-out flag.

The expected-outcome metadata turn each planted variant into an expected finding. They record the expected ACMG/AMP class and the classes accepted as correct (for example, likely pathogenic for a pathogenic variant), the expected report decision, the gene and condition, the criteria met in the eRepo curation, and whether the variant tests calling, interpretation, or both. The primary acceptance criterion is the two-tier report decision, and the five-tier class is secondary, because laboratories agree far more often on reportability than on the exact class (Mighton et al., 2022). Following the ACMG secondary-findings policy, P/LP variants in genes within the scope of the test are reported, whereas VUS, likely benign, and benign variants are not (Miller et al., 2021). UK guidance likewise excludes likely benign and benign variants from reports (Durkie et al., 2024). A laboratory with a different policy changes this rule in the suite configuration.

### 2.5 Genome construction and read simulation

Planted variants are merged into the germline background of the GIAB sample HG002 (Figure 2A), taken from the v4.2.1 small-variant benchmark on GRCh38 (Wagner et al., 2022a). The background gives the tertiary tool a realistic number of common germline variants, and any of them that reaches the report is scored as a false report. Background records that overlap a planted variant or lie within 500 bp of it are removed and logged, so each planted variant has an unambiguous haplotype. In somatic calling, false-positive calls were enriched near germline variants (Li et al., 2021), so the window could hide such errors. We therefore rebuilt one test set (chr17 and chr19) with a 10 bp window and otherwise identical settings (Supplementary Note S3). Because HG002 is male and the benchmark covers chromosomes 1–22 only (Wagner et al., 2022a), chromosome X is hemizygous outside the pseudoautosomal regions and has no background variants.

Paired-end reads are simulated with Mason (Holtgrewe, 2010) from the two haplotypes of the modified genome. In an independent comparison of six simulators against real NovaSeq 6000 data from yeast, Mason mimicked most features of modern Illumina data and produced exactly the requested number of reads. Its true alignments, like those of ART and NEAT, placed 97.1–98.3% of reads at their origin (Milhaven and Pfeifer, 2023). Its indel error rate, however, exceeded the expected rate by an order of magnitude (Milhaven and Pfeifer, 2023), so we compare simulated with real HG002 data before interpreting indel results (see Results).

A simulation profile sets read length, fragment size, coverage, and error profile. This study used 2 × 150-bp paired-end reads from fragments of 400 ± 60 bp (mean ± SD) at 30-fold coverage. The profile approximates the real HG002 genome used for comparison (2 × 151-bp reads, 30-fold coverage, mean insert size 409 bp), but its insert-size distribution is narrower (see Results). All Nordic clinical units target coverage above 30-fold for germline genomes (Lavrichenko et al., 2025). For fast regression runs, the framework also offers a region mode, which was not used in this study. It simulates reads only around each planted variant (1,500 bp on each side by default) and in further regions listed in a BED file. Paralogous loci, such as PMS2CL for PMS2 variants, belong in that file, because their reads keep the mapping ambiguity that difficult contexts are meant to test. Reads from the rest of the genome are still missing, so mapping is easier than in a whole genome, and the mode suits regression checks rather than estimates of absolute sensitivity. Only Mason is implemented; other simulators, such as ART (Huang et al., 2012), NEAT (Stephens et al., 2016), or Sandy (Miller et al., 2023b), would each need their own adapter. Because the choice of simulator can change the measured sensitivity of a caller (Yu et al., 2020), the simulator and its version are recorded with every result.

Before any system under test runs, the framework checks that every planted variant appears in Mason’s true alignments with the planted allele and adequate depth. AMP/CAP requires such verification of in silico samples (Roy et al., 2018), and insertion can fail silently: BAMSurgeon, for example, skips sites that fail its depth criteria and leaves them out of its truth (Ewing et al., 2015). A variant missing from the true alignments is a framework error (outcome class not_simulated) and is never counted against the pipeline.

### 2.6 Systems under test

Each system under test is wrapped by an adapter that declares its container image or executable, command line, inputs, and outputs. Secondary adapters take paired FASTQ files and return a BAM, a raw VCF, and a filtered VCF. Tertiary adapters take a VCF, and a tool-specific parser converts the output into canonical findings: position and alleles, gene, HGVS descriptions, assigned class, ACMG/AMP criteria, and report decision. Adding a tool therefore means writing an adapter and a parser, not changing the framework. If an adapter cannot supply an intermediate output, as with a vendor pipeline that emits only a filtered VCF, failure-stage attribution stops at the last stage it exposes.

A closed system that a laboratory cannot inspect, such as a commercial tool, is run through the same black-box adapter; DRAGEN, Franklin, and VarSome Clinical are named here only as examples of systems an adapter can wrap.

Secondary analysis in the reference implementation is nf-core/sarek (Garcia et al., 2020; Hanssen et al., 2024), release 3.10.0, run with Nextflow 26.10.2 (Di Tommaso et al., 2017). Reads were trimmed with fastp 1.1.0 (Chen et al., 2018), aligned with BWA-MEM2 2.2.1 (Vasimuddin et al., 2019), and called with DeepVariant 1.10.0 (Poplin et al., 2018). Reads were simulated from the no-alt analysis set, so none originates from an alternate contig; Sarek aligned them to its default GATK GRCh38 bundle, where 0.006% of the reads of one sample aligned to alternate contigs. Base quality score recalibration was skipped, because the DeepVariant developers advise against it: it slightly decreases the accuracy of DeepVariant (DeepVariant, 2026). Sarek already checks output checksums in continuous integration where possible and runs full-size tests on GIAB HG001 at each release, and its published germline benchmark measured calling only within GIAB high-confidence regions (Hanssen et al., 2024). The framework adds planted clinical variants, difficult contexts, and the report decision. It treats Sarek as a black box and reads the aligned reads (CRAM) and the variant calls. DeepVariant writes one VCF in which rejected candidates are marked in the FILTER column. Its PASS records therefore serve as the filtered call set.

DeepVariant can call chromosomes X and Y as haploid in male samples, with the pseudoautosomal regions (PAR) given in a BED file (DeepVariant, 2026). nf-core/sarek 3.10.0 passes neither option (its DeepVariant step receives no haploid contigs and an empty PAR file), so the baseline pipeline of this study called chromosome X of the male samples as diploid. Haploid calling with the GRCh38 PAR regions is the first controlled change of the regression test (see Results). We added both options through a Nextflow configuration file, without changing Sarek.

In this study, Sarek was run outside the framework, which adopted its outputs only after checking that the parameters, release, and input reads matched the pipeline configuration. Adopted outputs therefore carry the same provenance as runs that the framework starts itself.

Tertiary analysis uses BIAS-2015 (Eisenhart et al., 2025), version 3.0.0 with its data bundle of 1 March 2026. It is invoked as an external program and not redistributed. It is open source and automates 19 ACMG/AMP criteria with a rationale for each code. The nine remaining codes need patient or family information (Eisenhart et al., 2025), which a simulated genome cannot supply. The ClinVar-derived criteria PP5 and BP6 are switched on and off as an experimental factor. BIAS-2015 applies PP5 at stand-alone pathogenic strength to ClinVar variants with expert-panel or practice-guideline status (Eisenhart et al., 2025), and VCEP classifications are submitted to ClinVar (Preston et al., 2022). With PP5 enabled, the tool can therefore return the truth label of a curated variant directly, and comparing the two settings shows how much of the measured performance comes from database lookup. Switching them off does not remove all ClinVar evidence. In version 3.0.0, PS4 falls back on the number of ClinVar submitters with a P/LP assertion when no GWAS data are available (Bitscopic, 2026), and PS1 and PM5 use ClinVar variants at the same amino acid position (Eisenhart et al., 2025).

BIAS-2015 reads its evidence from a variant annotator, either Nirvana (Stromberg et al., 2017) or VEP (McLaren et al., 2016). We ran both, because a laboratory that changes its annotator changes the evidence its classifier receives. The annotator is therefore a second experimental factor. Nirvana 3.18.1 used Illumina’s annotation data, with ClinVar replaced by the February 2026 release and AlphaMissense (Cheng et al., 2023) added, as the BIAS-2015 documentation prescribes. VEP 115 used gnomAD v4.1 exome data (Chen et al., 2024), the ClinVar release of 1 February 2026, REVEL (Ioannidis et al., 2016), and AlphaMissense. Both annotators thus read the same ClinVar release, but population frequencies came from gnomAD v4.0 exomes in Nirvana and v4.1 in VEP.

The tertiary input was the PASS call set of the whole simulated genome, with multiallelic records split into biallelic ones. The VEP annotation that Sarek writes was not used. It lacks the custom annotations that BIAS-2015 requires and omits variants that are common in population databases. The report step of the tested pipeline reports every variant that BIAS-2015 classifies as P or LP, regardless of gene and zygosity. The report step belongs to the system under test. Its adapter can restrict reporting by class and by region; an inheritance rule, such as requiring two P/LP variants in a recessive gene, belongs to the laboratory’s own report step, which the adapter wraps. We added no such rule, so the measured specificity reflects the tested pipeline as configured.

### 2.7 Scoring and failure-stage attribution

The output of each run is scored against the truth at the VCF and report levels, and each failed expected finding is attributed to the first pipeline stage that went wrong.

At the VCF level, calls are compared with the truth after bcftools normalization, using hap.py with the vcfeval engine (Cleary et al., 2015; Krusche et al., 2019). A haplotype-aware engine is needed because many apparent differences between call sets are different representations of the same variant (Krusche et al., 2019). A match requires the correct genotype, the strictest of the three matching tiers (Krusche et al., 2019). We give no partial credit, which is inappropriate for small coding indels, where a slightly wrong call shifts the reading frame (Dunn and Narayanasamy, 2023). Results are stratified by variant type, indel length, and genomic context after the comparison, as recommended (Krusche et al., 2019). vcfdist (Dunn and Narayanasamy, 2023) or vcfeval alone can replace hap.py, which has not been updated since 2021.

Calls are scored in every simulated segment and in a window around each planted variant (50 bp plus the allele length on each side). Each segment is trimmed by 500 bp at both ends, because coverage falls near a segment end, and missed calls there would reflect the simulation rather than the pipeline. The window keeps a planted variant scored even when it lies in a trimmed end. The simulated genome is the reference plus the truth VCF, so the truth is complete wherever reads were simulated. Scoring therefore also covers regions that the GIAB benchmark excludes, where any call absent from the truth VCF is a false positive. It also covers chromosome X, which has no HG002 background, so every call there other than a planted variant is a false positive. The GIAB confident regions are reported as a separate stratum, so that simulated and real data can be compared on the same regions. Only PASS calls are scored; non-PASS calls are used only to attribute a missed variant to the filtering step.

At the report level, each canonical finding is matched to an expected finding by normalized position and alleles. The framework then checks whether the assigned class is acceptable and whether the report decision is the expected one. Reported findings that match no planted variant expected to be reported are recorded as false reports. Each false report is traced to its origin: a false-positive call, a true variant classified as reportable, a finding outside the scored regions, or a finding inside them without an exact allele match in the comparison output. A reported part of a planted variant that was called in another representation is linked to that variant and not counted again. Comparing a laboratory’s annotation with the correct one is often hard and may need manual checking (Duncavage et al., 2023). Fixing the expected outcome before the run makes this comparison automatic and repeatable and supports the required validation of automated annotation (Roy et al., 2018).

When an expected finding fails, the framework assigns the failure to the first stage that went wrong. It examines the intermediate outputs in pipeline order: Mason’s true alignments, the pipeline’s alignments, the raw and filtered VCF, the annotation, the classification, and the report (Figure 3A; outcome classes in Table 3). An upstream error is therefore never counted as an interpretation error, a concern that also motivates the layered attribution model of Mitev (2026). Automatic attribution also addresses a known weakness of end-to-end controls: when one control spans several steps, locating the error becomes the problem (Jennings et al., 2017).

**Figure 3.**
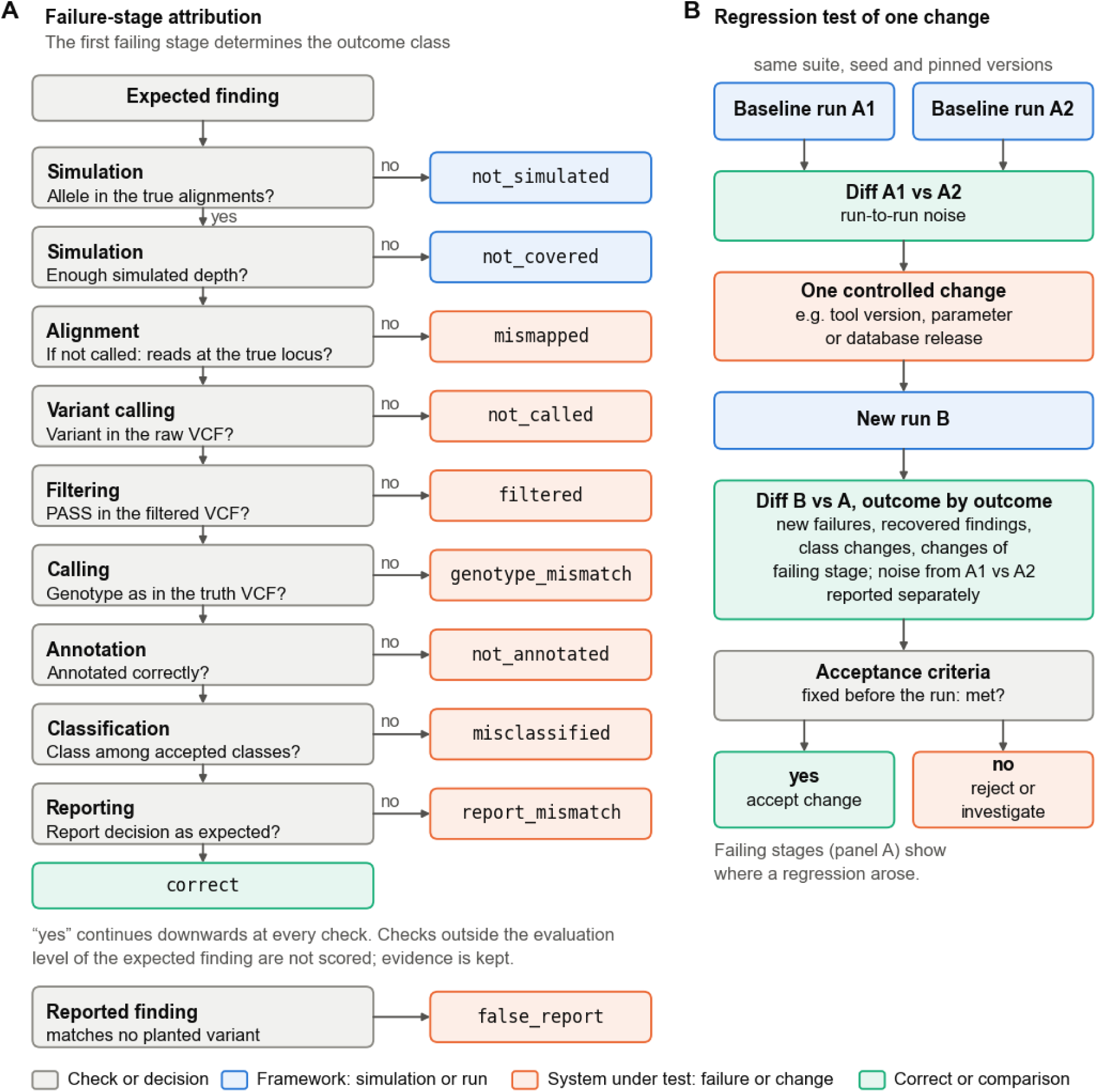
Failure-stage attribution and regression testing. (A) Checks applied to each expected finding in pipeline order. The first failing check determines the outcome class (Table 3); checks outside the evaluation level of the finding are not scored. (B) Regression test of one controlled change. Two baseline runs measure run-to-run noise, and the new run is compared with the baseline outcome by outcome against acceptance criteria fixed in advance.

The first stage that deviates is not always where a failure originates, because some failures arise from an interaction between stages. A variant in a low-coverage region may receive PM2, which guidelines advise against there (Durkie et al., 2024), and a population-frequency filter may remove a pathogenic variant with a high carrier frequency (Gargis et al., 2015). Such cases are assigned to the first stage whose output deviates from the expectation, and upstream evidence, such as depth at the locus, is kept in the outcome record. Each failure therefore has exactly one stage, but its record still shows the upstream conditions that contributed to it, so cross-stage dependencies can be analyzed (see Results).

### 2.8 Metrics and acceptance criteria

Calling performance is reported at two levels. For each planted variant, calling is part of its expected finding. Across all scored calls, it is reported as recall and precision per variant type and stratum (Krusche et al., 2019). Interpretation performance is reported on expected findings only (Figure 2B). Report-level sensitivity is the fraction of planted variants expected to be reported that are reported. Report-level specificity is the fraction of planted variants that must not be reported (B/LB variants, VUS, and single P/LP variants in recessive genes) that are correctly not reported; its denominator is known in advance. We do not report calling specificity, because the number of true negatives cannot be defined reliably (Krusche et al., 2019) and in silico data are better suited to false-negative than to false-positive rates (Duncavage et al., 2023). False reports are counted separately and kept out of the specificity estimate, because the number of variants that could be reported wrongly is not fixed. Five-tier concordance and per-criterion agreement with the eRepo curation are secondary measures.

All proportions carry 95% Wilson score intervals, which AMP/CAP suggests for accuracies close to 100% (Roy et al., 2018) and which matter most for small strata (Krusche et al., 2019). Report-level sensitivity and specificity are given by variant group (P/LP, B/LB, VUS, and recessive carriers) and by held-out status. Acceptance criteria are defined in a configuration separate from the test set and fixed before a run, because success criteria for pipeline changes must be defined in advance (Lavrichenko et al., 2025). The thresholds used here are an example configuration that shows how the criteria work. They are not proposed as clinical standards, and each laboratory sets its own. This study used five criteria: every planted variant called with the correct genotype, report-level sensitivity and specificity of at least 95%, a lower 95% interval bound of at least 80% for the sensitivity on held-out variants, and no false-positive call in the report. The 95% level is the reliability that AMP/CAP uses to size validation sets (Roy et al., 2018). We applied it to the point estimate, because with about 120 variants per group a lower interval bound of 95% would allow at most one error. For held-out variants, the criterion asks whether sensitivity drops markedly when the tool cannot draw on the data it was tuned on, so the interval must exclude a sensitivity below 80%. A regression test (Figure 3B) compares two runs outcome by outcome and lists every new failure, recovered finding, class change, and change of failing stage. Pipelines can produce minor differences in repeated runs (Lavrichenko et al., 2025). Where a repeated run of the baseline is available, it measures run-to-run noise, and differences already present between the two baseline runs are reported as noise rather than regression.

### 2.9 Implementation and availability

The framework is a Python package (Python 3.10 or later) with a command-line interface and YAML configuration of test suites. Data exchanged between steps follow versioned schemas, also published as JSON Schema. The package does not reimplement simulators, aligners, callers, or classifiers; it orchestrates them and implements variant selection, truth encoding, the expected-outcome model, finding-level comparison, failure-stage attribution, and the regression diff. External tools run from container images or as local executables; the provenance manifest records the image digest of every container. Nextflow and nf-core/sarek run on the host, and Sarek pulls its own per-process containers (Hanssen et al., 2024).

Every run writes a provenance manifest with the framework version and commit, SHA-256 checksums of all inputs and outputs, tool versions and container digests, database releases, the seed, and the wall time and peak memory of each step. This follows the recommendation that laboratories track software versions and assess each version change (Hume et al., 2019). Random decisions are derived from the suite seed by hashing, so repeated runs with the same inputs produce identical variant selections and truth files. The code is tested with continuous integration. The code is source-available under the Academic and Research License 1.0. Academic research, teaching and noncommercial research are free of charge, whereas clinical, diagnostic and commercial use, including the validation of a pipeline used in patient care, requires a written license from the author. The source code is available at https://github.com/ctu-fbme-dbi/insilico-clinval; release 1.0.0, which this article describes, is archived on Zenodo (Dvorský, 2026).

Runtime and cost were measured for the framework’s own steps, truth construction and read simulation, on Amazon Web Services (AWS). Secondary and tertiary analysis were not timed: they are systems under test, and their cost belongs to the laboratory’s pipeline. The comparison steps take seconds. We timed three test sets of increasing size, simulated from two chromosomes (chr17 and chr19), four chromosomes (chr12–chr15) and the whole genome, each with two samples at 30-fold coverage. Each repetition ran on a fresh on-demand m7i.2xlarge instance (8 vCPUs, 32 GiB memory, Stockholm region) with a data volume of fixed throughput. The two smaller sets were run three times and the whole-genome set twice. Inputs were read from Amazon S3 and checked against their SHA-256 checksums, and outputs were written back to S3. Wall time was recorded per step, and CPU, memory and disk use were sampled every 5 s; software setup and data transfer were timed separately. Cost was computed from the on-demand list prices of the instance and its volumes on 24 October 2026, and the deployment scripts are released with the code.

### 2.10 Ethics statement

Ethical approval was not required for this study in accordance with local legislation and institutional requirements, because no human participants were recruited and no new human data were generated. Publicly available reference data from the GIAB HG002 sample were used under their original consent.

## 3 Results

### 3.1 Fidelity and completeness of simulated data

A planted variant can test a pipeline only if it is present in the simulated reads, and AMP/CAP requires in silico samples to be verified (Roy et al., 2018). All 243 planted variants in ten samples of the five test sets were present in Mason’s true alignments on the intended haplotype (95% CI 98.4–100%; Figure 4A). Heterozygous variants had a median allele fraction of 0.50 (range 0.24–0.76), hemizygous and homozygous variants 1.0, and the four compound heterozygotes in *GAA* and *RPE65* were in trans, as planted. Reads carrying the planted allele had a median depth of 15 (minimum 3).

**Figure 4.**
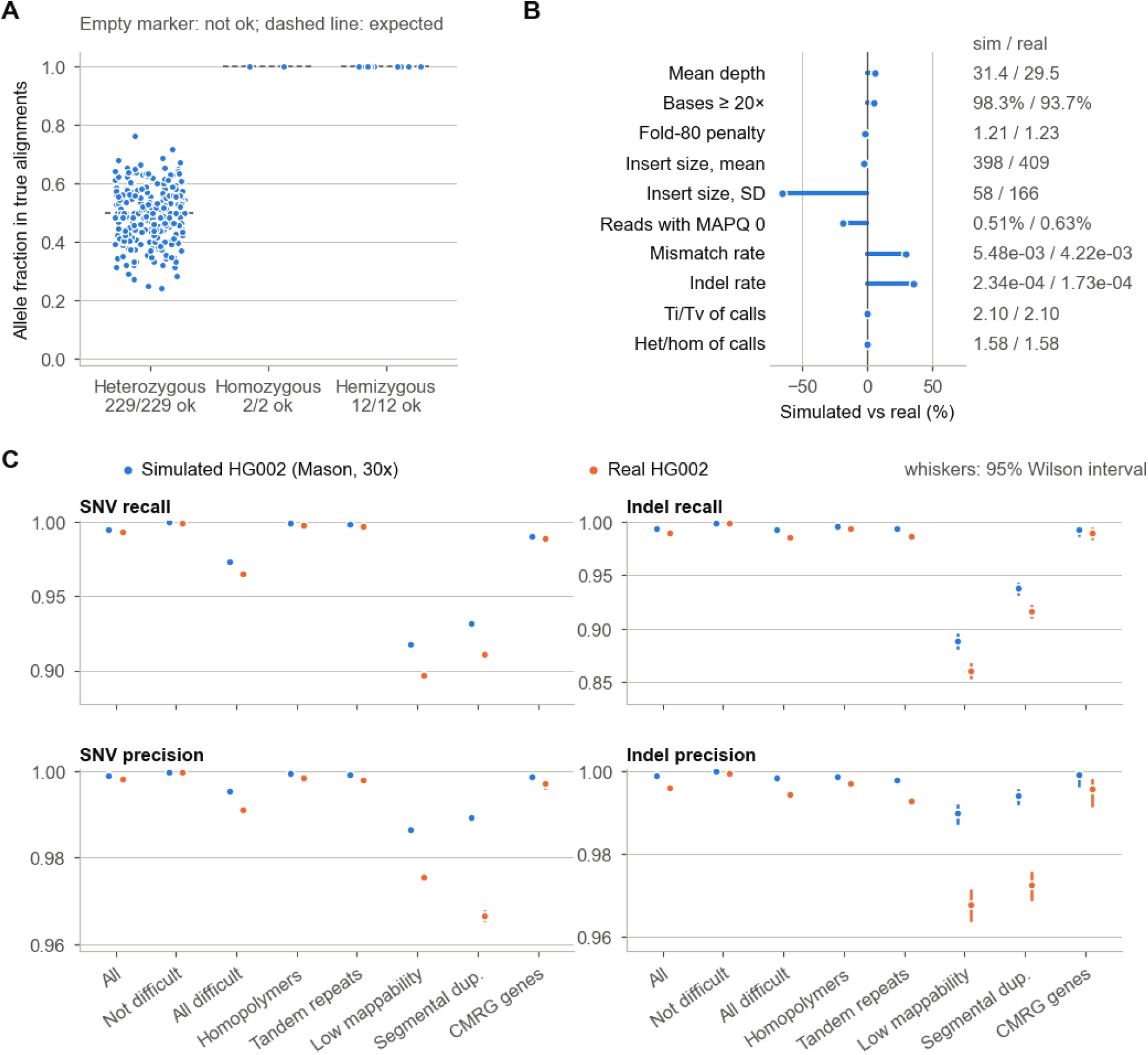
Completeness and fidelity of simulated data. (A) Allele fraction of each planted variant in Mason’s true alignments, by zygosity; dashed lines show the expected values. (B) Read-level metrics of simulated versus real HG002 data, as relative difference. (C) Recall and precision against the GIAB v4.2.1 benchmark by variant type and genome stratum; whiskers are 95% Wilson score intervals.

The check matters because planting can fail unnoticed. In a pilot at 3-fold coverage, 10 of 50 variants had no read with the planted allele, and the check marked the sample as incomplete.

To measure how realistic the simulated reads are, we processed the simulated sample and a real HG002 genome (NovaSeq 6000, PCR-free, 30-fold, 2 × 151 bp; public DeepVariant benchmarking data, see Data Availability Statement) with the same Sarek configuration. Coverage was similar (mean depth 31.4 and 29.5; 98.3% and 93.7% of bases at 20-fold or more), and mean insert sizes were close (398 and 409 bp), but the simulated insert sizes were narrower (standard deviation 58 and 166 bp), as set by the simulation profile (Figure 4B). Mismatch and indel rates against the reference were 30% and 35% higher in the simulated reads. Both rates include the true variants of HG002, so they compare the two datasets rather than measure sequencing error alone. The call sets had the same transition/transversion ratio (2.10) and heterozygous/homozygous ratio (1.58). Against the GIAB v4.2.1 benchmark, SNV recall was 99.5% on simulated and 99.3% on real data, and indel recall 99.4% and 98.9% (Figure 4C). Simulated recall was higher in every stratum. The gap was largest in low-mappability regions (indels 88.8% and 86.1%; SNVs 91.7% and 89.7%) and in segmental duplications (indels 93.7% and 91.6%).

The calling results in the following sections are therefore an upper bound, most of all in low-mappability regions and segmental duplications.

### 3.2 Secondary analysis performance

The four pipeline configurations share the outputs of Sarek, so their calling results are identical; they differ only in tertiary analysis. Secondary analysis called all 243 planted variants with the correct genotype (243/243; 95% Wilson interval 98.4–100%), 212 SNVs and 31 indels (Figure 5).

**Figure 5.**
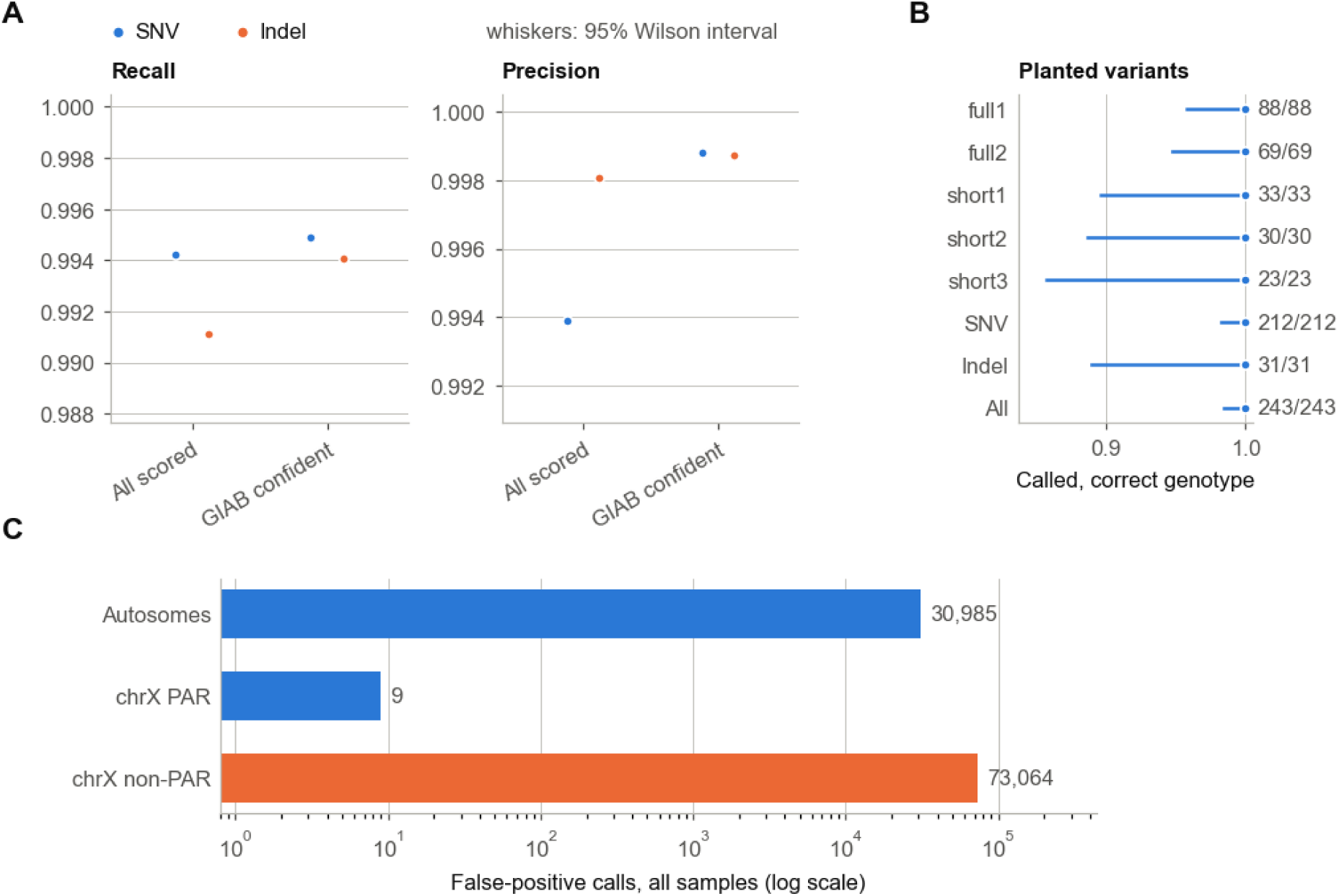
Calling performance of the secondary analysis. (A) Recall and precision by variant type in all scored regions of the ten samples and in the GIAB confident stratum. (B) Planted variants called with the correct genotype, by test set and by variant type. (C) False-positive calls by chromosome group, summed over all samples (log scale). Whiskers are 95% Wilson intervals. The four pipeline configurations share the Sarek outputs, so the panels hold for all of them.

Against the complete truth of the simulated segments of all ten samples, SNV recall was 99.4% and indel recall 99.1%, at a precision of 99.4% and 99.8%. Within the GIAB confident regions, recall was 99.5% for SNVs and 99.4% for indels, and precision was 99.9% for both.

False-positive calls were concentrated on chromosome X. The three test sets that simulate chromosome X had 73,064 false-positive calls on its non-PAR part, compared with 30,985 on all simulated autosomes together. Their origin is analyzed under failure-stage attribution below. As in any purely simulated test, these calling results are an upper bound for real data.

In the difficult-context test set, 273 of 292 planted variants were called with the correct genotype (93.5%; 95% CI 90.1–95.8%; Supplementary Table S1). All 240 ClinVar variants in segmental duplications, low-mappability regions, homopolymers, and tandem repeats were called correctly. The 19 failures concentrated in two contexts. Of 20 variants in regions that GRCh38 falsely duplicates, 15 failed: for 9 of them, reads carrying the allele were aligned away from the true locus, 5 were not called, and 1 was filtered. Of 5 variants in SMN1, 4 failed, 3 of them through mismapped reads; the PMS2 and OTOA variants were called. The framework assigned all 19 failures to alignment or calling, which is the expected weak point of short reads in these regions. At the report level, 14 of 15 curated P/LP variants were reported and all 10 B/LB variants were withheld with the baseline configuration.

The 500 bp background window did not change the results. In the control test set with a 10 bp window, 55 background records were restored near 18 of the 33 planted variants, 12 of them within one read length. Every planted variant had the same outcome under both windows in all four configurations (Supplementary Note S3, Supplementary Table S2). The control sample with negative variants had two additional false reports, both far from any planted variant.

In the indel-length experiment, every indel of 1–20 bp was called with the correct genotype (240/240; 95% CI 98.4–100%). Longer indels were missed more often: 71 of 74 indels of 21–35 bp were called (96%; 89–99%), 53 of 66 of 36–50 bp (80%; 69–88%), and 12 of 70 of 51–100 bp (17%; 10–28%; Supplementary Table S3). Between 36 and 50 bp, deletions were still mostly called (42/45) but insertions were not (6/15); above 50 bp, most deletions and insertions were missed (5/48 and 4/15).

### 3.3 Report-level performance of clinical interpretation

With the baseline configuration (Nirvana, PP5 and BP6 enabled), the pipeline reported 115 of 117 planted P/LP variants (98.3%; 95% CI 94.0–99.5%) and correctly withheld 106 of 126 planted variants that must not be reported (84.1%; 76.8–89.5%; Table 4). Specificity differed by group. B/LB variants were almost always withheld (84/86; 97.7%). In contrast, 13 of 34 VUS were upgraded and reported, and five of six carriers of a single P/LP variant in a recessive gene were reported.

**Table 4.** Report-level performance of the four pipeline configurations.

| **Measure** | **Nirvana** | **Nirvana, PP5/BP6 off** | **VEP** | **VEP, PP5/BP6 off** |
|---|---|---|---|---|
| Report sensitivity, P/LP (n = 117) | 115/117 = 98.3% (94.0–99.5) | 87/117 = 74.4% (65.8–81.4) | 115/117 = 98.3% (94.0–99.5) | 86/117 = 73.5% (64.9–80.7) |
| held out (n = 35) | 34/35 = 97.1% (85.5–99.5) | 23/35 = 65.7% (49.2–79.2) | 34/35 = 97.1% (85.5–99.5) | 26/35 = 74.3% (57.9–85.8) |
| not held out (n = 77) | 76/77 = 98.7% (93.0–99.8) | 61/77 = 79.2% (68.9–86.8) | 76/77 = 98.7% (93.0–99.8) | 58/77 = 75.3% (64.6–83.6) |
| Report specificity (n = 126) | 106/126 = 84.1% (76.8–89.5) | 110/126 = 87.3% (80.4–92.0) | 109/126 = 86.5% (79.5–91.4) | 111/126 = 88.1% (81.3–92.7) |
| B/LB (n = 86) | 84/86 = 97.7% (91.9–99.4) | 84/86 = 97.7% (91.9–99.4) | 86/86 = 100.0% (95.7–100.0) | 85/86 = 98.8% (93.7–99.8) |
| VUS (n = 34) | 21/34 = 61.8% (45.0–76.1) | 23/34 = 67.6% (50.8–80.9) | 23/34 = 67.6% (50.8–80.9) | 24/34 = 70.6% (53.8–83.2) |
| Recessive carriers (n = 6) | 1/6 = 16.7% (3.0–56.4) | 3/6 = 50.0% (18.8–81.2) | 0/6 = 0.0% (0.0–39.0) | 2/6 = 33.3% (9.7–70.0) |
| Acceptable class (n = 242) | 208/242 = 86.0% (81.0–89.8) | 165/242 = 68.2% (62.1–73.7) | 221/242 = 91.3% (87.1–94.3) | 158/242 = 65.3% (59.1–71.0) |
| False reports: false-positive calls | 33 | 32 | 49 | 48 |
| False reports: true background variants | 28 | 40 | 58 | 64 |
| False reports: planted negatives | 20 | 16 | 17 | 15 |

Values are k/n, percentage, and 95% Wilson interval over the five test sets (ten samples). Sensitivity: planted P/LP variants that must be reported and were reported; held out: absent from the BIAS-2015 benchmark of eRepo v2.2.0 (five P/LP variants of unknown status are in neither held-out row). Specificity: planted variants that must not be reported and were not. Acceptable class: planted variants classified within the accepted classes, among those with a class. False reports are counts; reported planted negatives also count against specificity. BP6, PP5: ClinVar-derived ACMG/AMP criteria; VEP, Ensembl Variant Effect Predictor; VUS, variant of uncertain significance.

The carriers were reported because BIAS-2015 classifies variants without regard to zygosity, and the pipeline reports every P/LP classification. We kept this property of the system under test. A reporting step that follows the secondary-findings policy would require two P/LP variants in a recessive gene (Miller et al., 2023a).

The reporting policy changed specificity but not the verdict. We rescored the runs with the rule that a recessive gene needs two P/LP variants. None of the reported carriers had a second reported P/LP variant in the same gene, so all were withheld. Specificity then rose from 84.1% to 88.1% (111/126; 95% CI 81.3–92.7%) with Nirvana and from 86.5% to 91.3% (115/126; 85.0–95.1%) with VEP. Both values stayed below the acceptance threshold of 95%.

Most upgrades of VUS did not depend on PP5. Six of the 13 upgraded VUS received PP5, but without PP5 and BP6 only two of them left the report. In 9 of the 13, BIAS-2015 applied at least one criterion that the VCEP had evaluated and recorded as not met, most often PS3 (five variants), which BIAS-2015 derives from literature mining (Eisenhart et al., 2025), and PP3 (four). Both truncating variants among them, in MYH7 and in the last exon of BRCA2, received PVS1, which their VCEPs had recorded as not met. Two further errors lay outside the classification rules. For the MYH7 variant, PP5 cited the ClinVar record of another allele at the same position (c.4588C>A). An OTC deletion received PVS1 because Nirvana annotated its left-aligned form as a splice-donor variant; in its 3′-shifted form, c.298+3_298+6del, the canonical splice site is intact, and with VEP the variant was correctly classified as VUS.

Report sensitivity depended on the ClinVar-derived criteria. Without PP5 and BP6, it fell from 98.3% to 74.4% (87/117; 65.8–81.4%) with Nirvana and to 73.5% (86/117; 64.9–80.7%) with VEP, while specificity rose slightly, to 87.3% and 88.1%. With Nirvana, 29 planted P/LP variants then received a class outside P/LP. This drop is larger than the one reported for an earlier version of BIAS-2015 on eRepo when PP5 alone was switched off, from 92.26% to 81.80% (Eisenhart et al., 2024).

Held-out variants, which are absent from the benchmark on which BIAS-2015 was tuned, kept their sensitivity when PP5 was enabled (34/35, 97.1%; others 76/77, 98.7%). Without PP5, sensitivity was lower for held-out variants with Nirvana (23/35, 65.7%; 49.2–79.2%) than for the others (61/77, 79.2%; 68.9–86.8%), but the intervals overlap, and with VEP the two groups were similar (74.3% and 75.3%). These data suggest, but do not show, an advantage of tuning on the curation set.

The annotator changed the assigned classes more than the report decisions for planted variants. Sensitivity was identical with both annotators (98.3%), and specificity was similar (84.1% with Nirvana, 86.5% with VEP). With VEP, more planted variants received an acceptable class (91.3% vs 86.0%), but more unplanned findings reached the report: 49 false-positive calls and 58 true background variants, against 33 and 28 with Nirvana.

As a secondary measure, the criteria assigned by BIAS-2015 agreed only partly with the expert curation. With Nirvana, the tool assigned 53.5% of the curated codes, and 40.0% of the codes it assigned were in the curation (codes compared without strength). Part of this gap is expected, because BIAS-2015 does not automate the codes that need patient or family information (Eisenhart et al., 2025).

Against the acceptance criteria (see Metrics and acceptance criteria), the baseline passed planted-variant calling (243/243), report sensitivity (98.3%, threshold 95%), and sensitivity for held-out variants (lower interval bound 85.5%, threshold 80%). It failed report specificity (84.1%, threshold 95%) and the criterion of no false-positive calls in the report (33, threshold 0). The specificity failure came from the 13 upgraded VUS and the 5 reported recessive carriers; B/LB variants alone met the threshold (84/86; 97.7%, 95% CI 91.9–99.4%). The carriers were reported because the report step reports every P/LP classification regardless of zygosity. Ten of the 33 false-positive calls came from the chromosome X artifact described below. Because the thresholds are an example, these failures show what the criteria detect rather than whether BIAS-2015 is fit for clinical use.

In the indel-length experiment, report decisions again depended on PP5. Of 85 P/LP indels in secondary-findings genes, 53 were reported with Nirvana and with VEP, 34 with Nirvana without PP5 and BP6, and 12 with VEP without them; all 35 B/LB indels were withheld in every configuration (Supplementary Table S3). Sixteen indels called in another representation were classified only in parts, as in the GAA case described below.

Synthetic novel variants tested interpretation without a database assertion (Supplementary Note S1). All 324 were called with the correct genotype, and with Nirvana BIAS-2015 assigned PVS1 and PM2 to all but one of them. Of the 324, 83 (25.6%; 95% CI 21.2–30.6%) were classified as P or LP and reported (Supplementary Table S4). BIAS-2015 classified PVS1 with PM2 at supporting strength as VUS and reached P or LP only when a further criterion applied, most often PM1 or PP3 (Supplementary Table S5). Splice-site variants were therefore reported most often (23/30), followed by nonsense (10/30) and frameshift variants (50/264). The criteria themselves were mostly as expected: 309 of 324 variants (95.4%; 92.5–97.2%) were classified P or LP or received a VUS based on PVS1 and PM2 alone, so the combining rule caused 226 of the 241 failures. Of the other 15, 14 had also received BP4, a computational prediction of a benign effect, and one did not receive PVS1. Switching off PP5 and BP6 changed the class of no synthetic variant, and the two annotators disagreed on one.

### 3.4 Failure-stage attribution and cross-stage dependencies

Over the five test sets, every planted variant passed the calling stages with the baseline configuration, so all 40 failures arose in interpretation (Figure 6). Thirty-four variants were misclassified (15 B/LB, 17 VUS, 1 P/LP, and 1 recessive carrier), 5 recessive carriers were classified correctly but reported, and 1 P/LP variant was not annotated. The reports also contained 82 findings outside the expected ones: 33 false-positive calls, 28 true background variants classified as P/LP, 20 planted negative controls, which also count against specificity, and 1 finding without an exact allele match.

**Figure 6.**
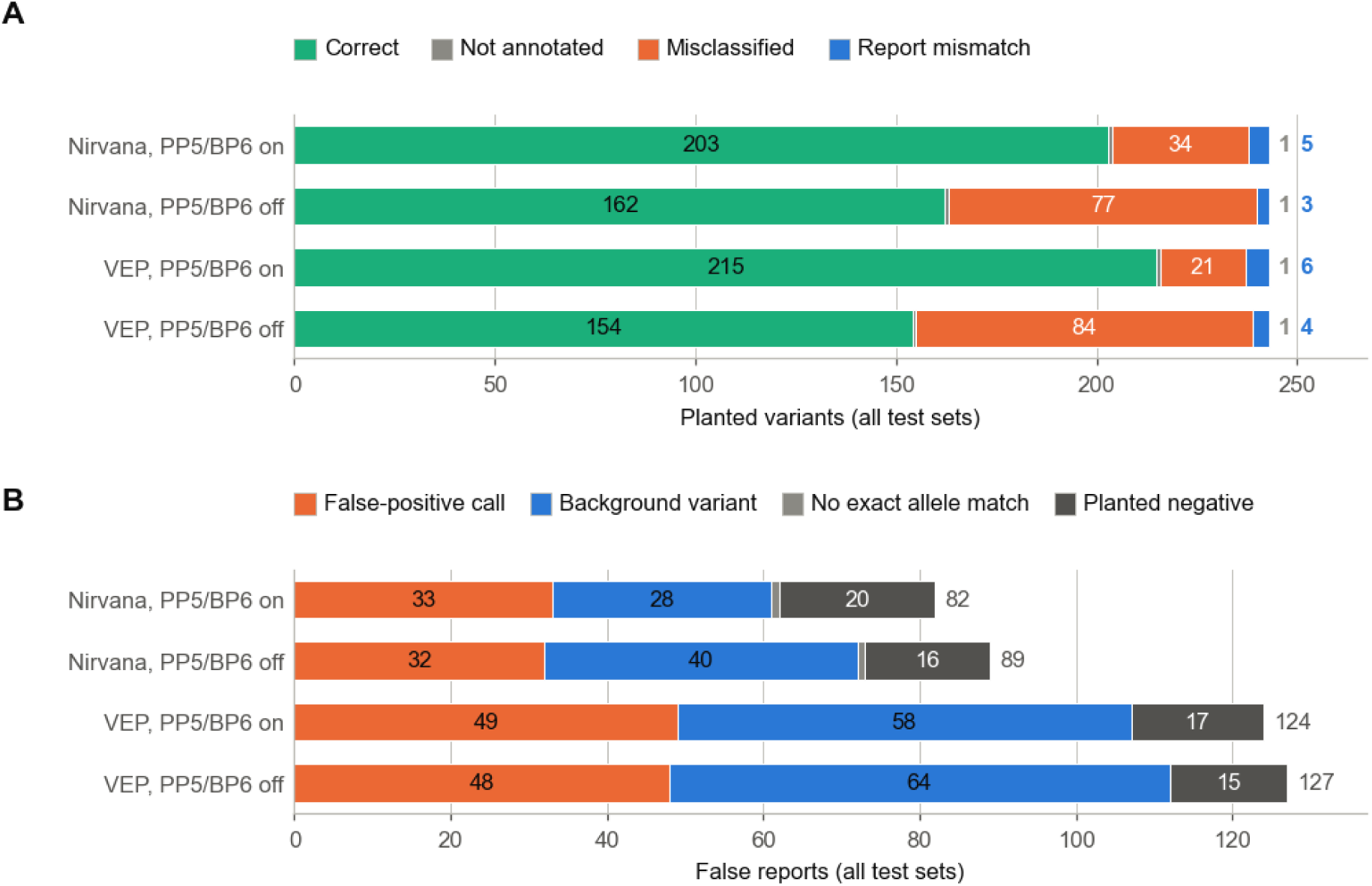
Failure-stage attribution of expected findings and origin of false reports, by pipeline configuration. (A) Outcome of the 243 planted variants; a failed variant carries the class of its first failing check (Table 3). (B) False reports by origin: false-positive call, true HG002 background variant, finding without an exact allele match in the comparison output, finding outside the scored regions, and reported planted negative control, which also counts against specificity.

One failure pattern dominated the false reports of the short3 test set (chromosomes 1, 10, and X). In its positive sample, DeepVariant made 12,184 PASS calls on the haploid part of chromosome X that matched no truth variant, and 12,176 of them were heterozygous. Their median allele fraction was 0.15 at a median depth of 13, so a typical call rested on two reads. These calls accounted for 10 of the 14 false reports of the two samples, and the framework assigned them to secondary analysis rather than interpretation.

To find their origin, we examined the 403 reads carrying the alternative allele at 200 randomly chosen sites. All came from the simulated haploid chromosome X, all had mapping quality 60, and all carried the same base in Mason’s true alignments. Their base quality was as high as that of reference bases (median 39). The calls therefore arose from simulated sequencing errors, not from mismapped reads. Two errors at the same position sufficed for a heterozygous call, which is consistent with chromosome X being called as diploid at half the autosomal coverage.

Real data also contain heterozygous calls on male chromosome X, but far fewer. The real HG002 genome, processed with the same pipeline, had 3,094 heterozygous calls on the non-PAR part of chromosome X, 524 of them with an allele fraction below 0.25. The simulated sample had 11,677 such low-fraction calls. Simulated reads therefore overstate this artifact (see Discussion).

Cross-stage dependencies also appeared between calling and annotation. A planted *GAA* variant, a deletion–insertion G>AT, was called as an insertion and an adjacent SNV. Together, the two calls form the correct haplotype, and hap.py scored the variant as found. BIAS-2015, however, classified each call separately and reported only the insertion as pathogenic, so the report named a variant other than the planted one. The framework recorded the planted variant as not annotated and linked the reported insertion to it through the hap.py records, so the failure was counted once.

### 3.5 Regression testing under controlled pipeline changes

As a first controlled change, we enabled haploid calling of chromosomes X and Y in DeepVariant and compared the new pipeline with the baseline on the short3 test set (Figure 7). Nothing else in the pipeline changed.

**Figure 7.**
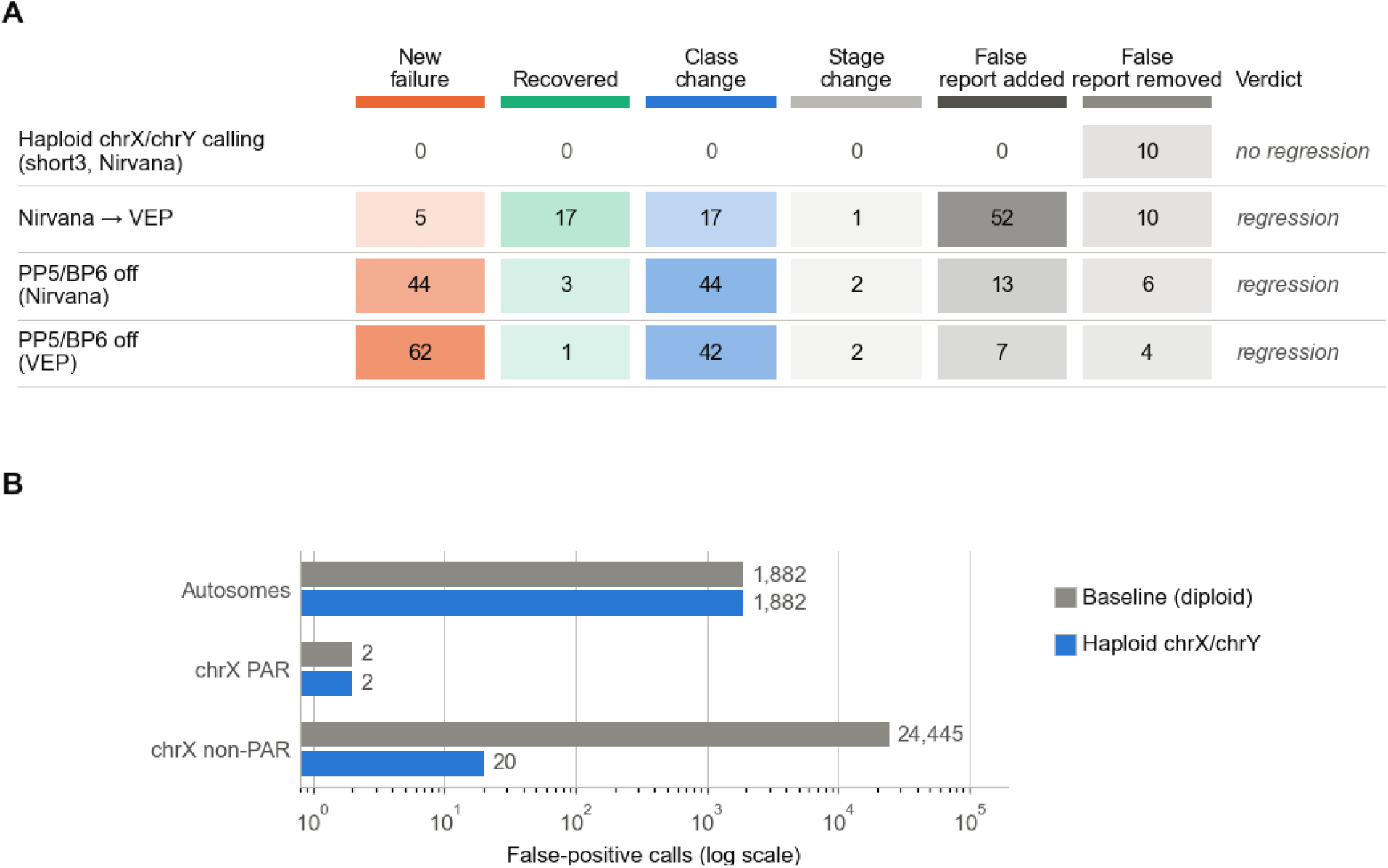
Regression testing of controlled pipeline changes. (A) Changes against the baseline for four controlled changes: haploid calling of chromosomes X and Y in DeepVariant (short3 test set), Nirvana replaced by VEP, and PP5/BP6 switched off with Nirvana and with VEP (all five test sets); cell shading scales with the count, and the last column gives the verdict of the comparison. (B) False-positive calls by chromosome group before and after haploid calling, both samples of the short3 test set (log scale).

The change removed the chromosome X artifact without side effects. All 23 planted variants had the same outcome as in the baseline. False reports fell from 6 to 1 in the positive sample and from 8 to 3 in the negative sample, and all 10 removed false reports were false-positive calls on chromosome X. False-positive calls on the non-PAR part of chromosome X fell from 12,182 to 9 and from 12,263 to 11, while recall of the planted variants there did not change. Autosomal calls were identical.

The new run realigned all reads, so for the autosomes it also served as a repeat of the baseline. All 527,488 and 527,496 PASS calls on chromosomes 1 and 10 of the two samples were identical between the runs, including genotypes. Run-to-run noise of this pipeline was therefore zero on this test set, and every difference reported above is attributable to the change.

The same comparison was applied to two changes of the tertiary analysis on all five test sets, each with its own baseline. Calling was identical in all of them, because the pipelines shared the Sarek outputs, so every difference arose in interpretation or reporting. The baseline with BIAS-2015 and Nirvana was run twice on the same Sarek outputs. Both runs gave the same outcome for every planted variant and the same false reports in all five test sets, so run-to-run noise of the tertiary analysis was zero.

Replacing Nirvana with VEP changed the outcome of 40 planted variants. Seventeen failures were recovered, mostly B/LB variants that Nirvana had classified as VUS (11) and VEP as benign. Five new failures appeared, and 17 variants changed class without changing the report decision, most often from likely pathogenic to pathogenic (12). The report itself changed more: 52 unplanned findings were added and 10 removed. Of the added findings, 34 were real background variants classified as P/LP and 18 were false-positive calls. Report-level sensitivity was the same with both annotators (Table 4), so a laboratory that judged the change only by sensitivity would miss the extra findings in its reports.

Switching off PP5 and BP6 produced the largest set of changes. With Nirvana, 44 planted variants failed that had passed before: 27 reportable variants were downgraded to VUS and left the report, 16 B/LB variants became VUS, and one pathogenic variant became likely benign. A further 35 pathogenic variants stayed in the report as likely pathogenic, and only three failures were recovered. With VEP, the same change produced 62 new failures: 29 reportable variants were downgraded to VUS, 32 B/LB variants became VUS, and one benign variant was classified as likely pathogenic and reported. These changes are the variant-level view of the sensitivity loss in Table 4. In every comparison the diff assigned each change to its stage and listed the affected variants, which is the information a laboratory needs to decide whether to accept a change.

### 3.6 Runtime and cost

Simulation time grew linearly with the number of simulated read pairs (7.4 min per 10 million pairs, R² = 0.999 over 16 samples). One 30-fold sample took 9.5 ± 0.6 min for two chromosomes (14.2 million read pairs), 34.0 ± 0.6 min for four chromosomes (45.7 million) and 3.7 ± 0.1 h for the whole genome (298 million; mean ± SD; Table 5). Mason took 61% of this time in the chromosome sets and 78% in the whole genome, where it overlapped with the conversion of its true alignments; BAM sorting took 9% and 16%. The eight vCPUs were 60–66% busy on average. Peak memory was 7.6–7.8 GB in every set, so memory did not grow with the size of the set, whereas disk use did: the whole-genome runs peaked at 447 GB. All repetitions produced byte-identical FASTQ files, and every planted variant was present in every sample.

**Table 5.** Runtime and cost of simulating the test sets on AWS.

| **Measure** | **Two chromosomes (chr17, chr19)** | **Four chromosomes (chr12–chr15)** | **Whole genome** |
|---|---|---|---|
| Runs / samples | 3 / 6 | 3 / 6 | 2 / 4 |
| Read pairs per sample (million) | 14.2 | 45.7 | 298.2 |
| Simulation per sample (min) | 9.5 ± 0.6 | 34.0 ± 0.6 | 220.5 ± 4.9 |
| of which Mason (min) | 5.8 ± 0.4 | 20.8 ± 0.4 | 171.6 ± 3.3 |
| CPU core-hours per sample | 0.83 ± 0.06 | 2.71 ± 0.07 | 18.97 ± 0.58 |
| Peak memory (GB) | 7.6 | 7.7 | 7.8 |
| Peak disk, maximum (GB) | 60 | 162 | 447 |
| FASTQ files per sample (GiB) | 3.5 | 11.2 | 73.0 |
| Test set: truth and two samples (min) | 21.5 ± 1.6 | 70.5 ± 1.3 | 443.4 ± 7.9 |
| Cost per sample (USD) | 0.08 | 0.28 | 2.14 |
| Cost per test set (USD) | 0.17 | 0.58 | 4.31 |
| Storage of the test set in Amazon S3 (USD per month) | 0.29 | 0.95 | 6.18 |

Mean ± SD over repetitions; per-sample values pool both samples of a test set. Each repetition ran on a fresh on-demand m7i.2xlarge instance (8 vCPUs, 32 GiB memory, AWS Stockholm region). Peak disk is the maximum over repetitions. Costs are on-demand list prices of 24 October 2026. Secondary and tertiary analysis are not included. AWS, Amazon Web Services; S3, Amazon Simple Storage Service.

At on-demand list prices, one sample cost 0.08, 0.28 and 2.14 USD for the three sets, and the whole-genome test set of two samples 4.31 USD. For comparison, secondary analysis of one normal whole genome cost 20.82 USD with Sarek 3.1.1 on AWS (Hanssen et al., 2024); that figure comes from a different setup and serves only as an order of magnitude. A test set is simulated once and reused for every pipeline release, so its cost recurs only when the variant selection or the simulation profile changes; keeping the whole-genome set in Amazon S3 costs 6.18 USD per month. Machine setup and input transfer added under 3 min per run, and uploading the whole-genome outputs 15 min; a prebuilt machine image would remove most of the setup. Several clinical units already use reduced data sets, such as selected chromosomes, for quick complete pipeline runs, because a whole-genome pipeline can take days of CPU time (Lavrichenko et al., 2025). The two-chromosome set (21.5 min and 0.17 USD per test set) fits this use for frequent regression runs.

## 4 Discussion

The framework tested a clinical pipeline end to end against a known truth and assigned every failure to a stage. Three findings stand out. First, secondary analysis found every planted variant, so all failures of planted variants arose in interpretation; the largest was a dependence on ClinVar-derived criteria, without which report sensitivity fell by 24 percentage points. Second, report specificity was limited by VUS upgraded to P/LP and by carriers in recessive genes, whereas B/LB variants were almost always withheld. Third, false-positive calls produced about half of the unplanned findings in the reports. In one test set, they came from an artifact on the haploid chromosome X, which the framework traced to diploid calling of simulated sequencing errors and which one controlled change removed without side effects.

Other approaches handle planting failures differently: read editing with insiM placed 7 of 49 variants at the wrong position (Roy et al., 2025), and Eduomics kept only the 38.2% of injected variants that its own workflow called (Sola et al., 2026). Because the framework checks the true alignments rather than variant calls, a variant that was never simulated is never counted as a pipeline error.

Simulated reads cannot model all artifacts of wet-laboratory sequencing, and purely simulated data tend to overestimate performance, especially in repetitive regions (Li et al., 2021; Duncavage et al., 2023). The calling results are therefore an upper bound. The comparison with real HG002 data (Figure 4) shows where the gap is largest, and results in those strata need cautious interpretation. The choice of simulator can also change the measured sensitivity of a caller (Yu et al., 2020), so the simulator is recorded with every result. Mason is also known to overstate the indel error rate (Milhaven and Pfeifer, 2023), which likely explains part of the higher indel rate of the simulated reads. Base quality may be one cause of this gap. Mason gave its sequencing errors the same base quality as correct bases, whereas in tumor data built by editing real sequencing reads, false-positive somatic SNV calls were enriched at low base quality (Li et al., 2021). This difference likely contributed to the roughly 20-fold excess of low-fraction heterozygous calls on simulated chromosome X compared with real HG002. The direction of this bias determines which results it affects. Sequencing errors with high base quality make false-positive calls more likely, so specificity measured on simulated reads is conservative, whereas sensitivity remains an upper bound. Callers and filters that weigh base quality may be penalized more than on real data, so comparisons of such tools need a simulator whose errors follow the base quality. ART draws substitution errors from empirical position-dependent quality profiles (Huang et al., 2012), and NEAT inserts errors with a probability tied to quality scores learned from real data (Stephens et al., 2016); ART, however, underestimates the indel error rate by an order of magnitude (Milhaven and Pfeifer, 2023). Mason sets the quality of erroneous bases separately from that of correct bases, so a lower value is a change of the simulation profile. Because the framework compares simulated with real data, the choice of simulator and profile can be checked before a test set is used.

Whether the upgrades of VUS are specific to BIAS-2015 cannot be answered with one tool. Upgrades of VUS are, however, a known tendency of automated classifiers: three of four broad-spectrum tools classified more than half of ClinGen VUS as P or LP (Costa et al., 2025). In our data, most upgrades rested on criteria that the expert panels had evaluated and rejected, which suggests that generic rules, rather than missing evidence, caused them. Any classifier that does not apply the gene-specific rules of the VCEPs may share this weakness, and the framework can measure it for each one.

Interpretation results carry two further limits. First, the expected classes come from ClinGen eRepo, on which BIAS-2015 was tuned (Eisenhart et al., 2025), and its ClinVar criteria can return the curated label directly. Measured performance may therefore be optimistic. The framework offers three mitigations: switching PP5 and BP6 off, flagging held-out variants, and planting synthetic variants that no database classifies. In this study, the first two were applied to all test sets and the third in a supplementary experiment. Switching PP5 and BP6 off does not remove the effect completely, because PS4, PS1, and PM5 also draw on ClinVar (see Systems under test). Without PP5 and BP6, 72 of the 87 reported P/LP variants still received PS4 from ClinVar submitter counts. This configuration is therefore an upper estimate of performance without database lookup, and only synthetic variants test the classification logic alone. It also follows the ClinGen SVI recommendation that laboratories stop using PP5 and BP6 (Biesecker et al., 2018). Without a database assertion, BIAS-2015 classified about three quarters of the synthetic loss-of-function variants as VUS. The failure arose where two recommendations meet. BIAS-2015 applied PM2 at supporting strength, as the ClinGen SVI recommends, but combined the criteria by the rules of 2015, under which PVS1 with one supporting criterion is VUS (Richards et al., 2015). The same SVI recommendation adds a rule that classifies this combination as likely pathogenic (ClinGen Sequence Variant Interpretation Working Group, 2020), and under the ACGS guidelines, which keep PM2 at moderate strength, these variants would be likely pathogenic or pathogenic (Durkie et al., 2024). A tool that adopts an updated criterion without the matching combining rule thus departs from both recommendations, so laboratories need to state which combining rules their pipeline follows. The framework thus located the failure in the combining rule rather than in the criteria, a gap that curated variants can hide while PP5 supplies the class (Supplementary Note S1). The held-out flag rests on the public benchmark file of BIAS-2015, which is not an exact copy of the eRepo snapshot: many of its indel alleles are malformed, and six planted variants therefore have an unknown status. Second, criteria that need patient or family information, such as PS2, PM3, PP1, PP4, and BS4, are not automated by BIAS-2015 (Eisenhart et al., 2025). A single simulated genome cannot test them.

The scope is also limited. The results describe the tested pipeline on curated variants in 29 genes of the ACMG secondary-findings list, not variant interpretation in general. These genes were included because eRepo held expert curations for them, and their VCEPs have written gene-specific rules. Genes with less evidence in ClinVar or without such rules may give lower sensitivity and other failure patterns, because report sensitivity depended on ClinVar-derived criteria. The framework itself does not depend on the gene set. The reference implementation covers SNVs and indels of 1–20 bp. Copy-number and structural variants, repeat expansions, and mosaicism are out of scope. The planted indels of the five test sets were 1–5 bp long. In a supplementary experiment, calling was complete up to 20 bp and fell to 80% at 36–50 bp and 17% at 51–100 bp, which supports the scope of 1–20 bp for this pipeline (Supplementary Note S2). Report-level specificity is measured only on planted B/LB variants, VUS, and recessive carriers. Other wrongly reported variants are counted as false reports, which have no fixed denominator. Three categories fall below 59 variants, the minimum that AMP/CAP sets per variant type for 95% reliability at 95% confidence (Roy et al., 2018) and that we use as the target per category: VUS (34), held-out P/LP variants (36), and carriers in recessive genes (6). Their estimates show how the framework reports such strata but do not demonstrate reliability. A laboratory that uses the framework for formal validation must enlarge these categories. The eRepo export used here holds several hundred further curations of each kind in the same genes, so this needs only larger selection counts. The HG002 benchmark has no truth on chromosome X (Wagner et al., 2022a). The simulated chromosome X therefore carries planted variants only, which tests haploid calling but not calling among background variants. Background variants within 500 bp of a planted variant were removed. In a control with a 10 bp window, no planted variant changed its outcome, which excludes a large effect of the window but not a small one; the control covered two chromosomes and did not test two variants in the same codon. Finally, the framework supplements validation with human samples but does not replace it (Roy et al., 2018; Duncavage et al., 2023).

The chromosome X artifact also has a practical side. With its default settings, nf-core/sarek 3.10.0 calls chromosome X of male samples as diploid, and on the real HG002 genome this produced 3,094 heterozygous calls in a region where a male genome carries one copy. Laboratories that use this pipeline for male samples may want to enable haploid calling and confirm the effect with a regression test such as ours.

The framework is not limited to validating one pipeline. A test set is defined in configuration, so a laboratory can build new sets for its own genes and variant types. The simulation profile sets coverage, read length, and fragment size, so the same set can show how a pipeline performs under different sequencing conditions. Because the truth is fixed and components are exchanged through adapters, the framework can also compare secondary pipelines, aligners, or variant callers on the same simulated reads, and its comparison with real data could compare read simulators. None of these uses was tested in this study, which tested one secondary and one tertiary system. It therefore shows that the framework works end to end, not that every adapter works. Each of the four controlled changes ran through its own adapter configuration, while the truth, the comparison, and the attribution stayed the same, so a further pipeline needs a new adapter but no new evaluation logic. A short test set is simulated in about 20 min (Table 5) and can then be reused, so such comparisons are inexpensive.

Several extensions follow from these limitations. Simulated trios would make de novo and in-trans evidence (PS2, PM3) testable. Copy-number and structural variants and long reads would extend the scope beyond small variants. Copy-number and structural variants would change three steps. The truth would encode them with symbolic alleles and breakpoints, and the simulated haplotypes would carry the rearranged sequence. Comparison would need matching of breakpoints, size, and inserted sequence, which small-variant engines such as hap.py and vcfeval do not provide (Krusche et al., 2019). Interpretation would follow the points-based ACMG/ClinGen standard for copy-number variants (Riggs et al., 2020). The expected-outcome model and failure-stage attribution would remain unchanged. Running the framework in continuous integration would turn each pipeline change into an automatic regression test. Sarek already runs full-size tests on the GIAB sample HG001 at each release (Hanssen et al., 2024). Shared in silico test sets could also support external quality assessment of interpretation, not only of detection. Simulating systematic sequencing artifacts would narrow the realism gap described above.

In conclusion, the framework extends truth-based validation from variant calls to the clinical report. Because the expected variants, classes, and report decisions are fixed before any reads are simulated, it measures report-level sensitivity and specificity, which studies on real samples without a designed truth cannot measure directly, and it attributes each failure to the stage where it first appears. In this study, this showed that the weak point of the tested pipeline was interpretation rather than calling, and it traced an unexpected source of false reports to a default setting that one controlled change removed. Laboratories can use the framework both to validate a pipeline before clinical use and to rerun the same tests after every change, so that a regression is detected before it reaches a patient report.

## 6 Author Contributions

BD: Conceptualization, Methodology, Software, Validation, Formal analysis, Investigation, Resources, Data curation, Writing – original draft, Writing – review & editing, Visualization, Project administration. MK: Funding acquisition, Validation, Writing – review & editing. KK: Validation, Writing – review & editing. OK: Validation, Writing – review & editing. RK: Supervision, Validation, Writing – review & editing.

## 8 Generative AI statement

The authors declare that generative AI was used in the creation of this manuscript. Claude Opus 5.5 (Anthropic) was used to draft and edit text, check citations and write analysis code; the authors verified all output and take full responsibility for the content.

## 10 Supplementary Material

Supplementary Note S1: synthetic novel variants. Supplementary Note S2: indel-length experiment. Supplementary Note S3: background-window control. Supplementary Table S1: outcomes of the difficult-context test set by selection rule and pipeline configuration. Supplementary Table S2: outcome of each planted variant with a 500 bp and a 10 bp background window. Supplementary Table S3: calling and report decisions by indel length. Supplementary Table S4: synthetic variants by type, scored with two expected classes. Supplementary Table S5: ACMG/AMP criteria assigned by BIAS-2015 to the synthetic variants.

## 11 Data Availability Statement

The simulated datasets, variant manifests and expected-outcome metadata generated for this study are available on Zenodo (Dvorský et al., 2026; https://doi.org/10.5281/zenodo.23863823). The source code of the framework is available at https://github.com/ctu-fbme-dbi/insilico-clinval as source-available code under the Academic and Research License 1.0 (see Implementation and availability); the version described in this article is archived at https://doi.org/10.5281/zenodo.23859473 (Dvorský, 2026).

The real HG002 reads used for the fidelity comparison (Illumina NovaSeq 6000, PCR-free, 30-fold, 2 × 151 bp) are public DeepVariant benchmarking data (gs://deepvariant/benchmarking/fastq/wgs_pcr_free/30x/).

## References

Bastida-Lertxundi, N., Martí-Carrera, I., Laña-Ruíz, B., Martínez-Múgica Barbosa, O., Muguerza-Iraola, R., Sáez-Villaverde, R., et al. (2025). Evaluating seven bioinformatics platforms for tertiary analysis of genomic data from whole exome sequencing in a pilot group of patients. Adv. Lab. Med. 6, 28–36. doi: 10.1515/almed-2025-0031

Biesecker, L. G., Harrison, S. M., and ClinGen Sequence Variant Interpretation Working Group (2018). The ACMG/AMP reputable source criteria for the interpretation of sequence variants. Genet. Med. 20, 1687–1688. doi: 10.1038/gim.2018.42

Bitscopic (2026). BIAS-2015 changelog, version 3.0.0. Available at: https://github.com/bitscopic/BIAS-2015/blob/v3.0.0/CHANGELOG.md (accessed 1 November 2026).

Behera, S., LeFaive, J., Orchard, P., Mahmoud, M., Paulin, L. F., Farek, J., et al. (2023). FixItFelix: improving genomic analysis by fixing reference errors. Genome Biol. 24, 31. doi: 10.1186/s13059-023-02863-7

Chen, S., Zhou, Y., Chen, Y., and Gu, J. (2018). fastp: an ultra-fast all-in-one FASTQ preprocessor. Bioinformatics 34, i884–i890. doi: 10.1093/bioinformatics/bty560

Chen, S., Francioli, L. C., Goodrich, J. K., Collins, R. L., Kanai, M., Wang, Q., et al. (2024). A genomic mutational constraint map using variation in 76,156 human genomes. Nature 625, 92–100. doi: 10.1038/s41586-023-06045-0

Cheng, J., Novati, G., Pan, J., Bycroft, C., Žemgulytė, A., Applebaum, T., et al. (2023). Accurate proteome-wide missense variant effect prediction with AlphaMissense. Science 381, eadg7492. doi: 10.1126/science.adg7492

Cleary, J. G., Braithwaite, R., Gaastra, K., Hilbush, B. S., Inglis, S., Irvine, S. A., et al. (2015). Comparing variant call files for performance benchmarking of next-generation sequencing variant calling pipelines. bioRxiv [Preprint]. doi: 10.1101/023754

ClinGen Sequence Variant Interpretation Working Group (2020). ClinGen Sequence Variant Interpretation recommendation for PM2 – Version 1.0 (approved 4 September 2020). Available at: https://www.clinicalgenome.org/site/assets/files/5182/pm2_-_svi_recommendation_-_approved_sept2020.pdf (accessed 31 October 2026).

Costa, M., García S., A., León, A., and Pastor, O. (2025). The promises and pitfalls of automated variant interpretation: a comprehensive review. Brief. Bioinform. 26, bbaf545. doi: 10.1093/bib/bbaf545

Danecek, P., Bonfield, J. K., Liddle, J., Marshall, J., Ohan, V., Pollard, M. O., et al. (2021). Twelve years of SAMtools and BCFtools. GigaScience 10, giab008. doi: 10.1093/gigascience/giab008

DeepVariant (2026). DeepVariant 1.10 documentation: README and DeepVariant details (r1.10 branch). Available at: https://github.com/google/deepvariant/tree/r1.10 (accessed 30 October 2026).

Di Tommaso, P., Chatzou, M., Floden, E. W., Barja, P. P., Palumbo, E., and Notredame, C. (2017). Nextflow enables reproducible computational workflows. Nat. Biotechnol. 35, 316–319. doi: 10.1038/nbt.3820

Duncavage, E. J., Abel, H. J., Merker, J. D., Bodner, J. B., Zhao, Q., Voelkerding, K. V., et al. (2016). A model study of in silico proficiency testing for clinical next-generation sequencing. Arch. Pathol. Lab. Med. 140, 1085–1091. doi: 10.5858/arpa.2016-0194-CP

Duncavage, E. J., Coleman, J. F., de Baca, M. E., Kadri, S., Leon, A., Routbort, M., et al. (2023). Recommendations for the use of in silico approaches for next-generation sequencing bioinformatic pipeline validation: a joint report of the Association for Molecular Pathology, Association for Pathology Informatics, and College of American Pathologists. J. Mol. Diagn. 25, 3–16. doi: 10.1016/j.jmoldx.2022.09.007

Dunn, T., and Narayanasamy, S. (2023). vcfdist: accurately benchmarking phased small variant calls in human genomes. Nat. Commun. 14, 8149. doi: 10.1038/s41467-023-43876-x

Durkie, M., Cassidy, E. J., Berry, I., Owens, M., Turnbull, C., Scott, R. H., et al. (2024). ACGS best practice guidelines for variant classification in rare disease 2024. Version 1.2. Association for Clinical Genomic Science. Available at: https://www.acgs.uk.com/media/12533/_media_12533_uk-practice-guidelines-for-variant-classification-v12-2024.pdf (accessed 31 October 2026).

Dvorský, B. (2026). insilico-clinval: end-to-end in silico validation and regression testing of clinical genome sequencing pipelines (version 1.0.0) [Software]. Zenodo. doi: 10.5281/zenodo.23859473

Dvorský, B., Komárková, M., Klempíř, O., Kolinová, K., and Krupička, R. (2026). Simulated test sets and evaluation results for in silico validation of a clinical genome sequencing pipeline (version 1.0) [Dataset]. Zenodo. doi: 10.5281/zenodo.23863823

Dwarshuis, N., Kalra, D., McDaniel, J., Sanio, P., Alvarez Jerez, P., Jadhav, B., et al. (2024). The GIAB genomic stratifications resource for human reference genomes. Nat. Commun. 15, 9029. doi: 10.1038/s41467-024-53260-y

Eisenhart, C., Mewton, J., Brickey, R., and Bayat, V. (2024). Automating ACMG variant classifications using BIAS-2015: an algorithm overview and benchmark against the FDA-approved eRepo dataset. medRxiv [Preprint]. doi: 10.1101/2024.09.18.24313916

Eisenhart, C., Brickey, R., Nadon, B., Mewton, J., and Bayat, V. (2025). Automating ACMG variant classifications with BIAS-2015 v2.1.1: algorithm analysis and benchmark against the FDA-approved eRepo dataset. Genome Med. 17, 148. doi: 10.1186/s13073-025-01581-y

Ewing, A. D., Houlahan, K. E., Hu, Y., Ellrott, K., Caloian, C., Yamaguchi, T. N., et al. (2015). Combining tumor genome simulation with crowdsourcing to benchmark somatic single-nucleotide-variant detection. Nat. Methods 12, 623–630. doi: 10.1038/nmeth.3407

Furtado, L. V. (2024). In silico options for assay validation. J. Appl. Lab. Med. 9, 180–182. doi: 10.1093/jalm/jfad099

Garcia, M., Juhos, S., Larsson, M., Olason, P. I., Martin, M., Eisfeldt, J., et al. (2020). Sarek: a portable workflow for whole-genome sequencing analysis of germline and somatic variants. F1000Research 9, 63. doi: 10.12688/f1000research.16665.2

Gargis, A. S., Kalman, L., Bick, D. P., da Silva, C., Dimmock, D. P., Funke, B. H., et al. (2015). Good laboratory practice for clinical next-generation sequencing informatics pipelines. Nat. Biotechnol. 33, 689–693 (supplementary information). doi: 10.1038/nbt.3237

Hansen, N. F., Dwarshuis, N., Ji, H. J., Rhie, A., Loucks, H., Logsdon, G. A., et al. (2026). A complete diploid human genome benchmark for personalized genomics. Cell 189, 4857–4875.e31. doi: 10.1016/j.cell.2026.06.016

Hanssen, F., Garcia, M. U., Folkersen, L., Pedersen, A. S., Lescai, F., Jodoin, S., et al. (2024). Scalable and efficient DNA sequencing analysis on different compute infrastructures aiding variant discovery. NAR Genom. Bioinform. 6, lqae031. doi: 10.1093/nargab/lqae031

Holtgrewe, M. (2010). Mason: a read simulator for second generation sequencing data. Technical Report B-10-06. Berlin: Fachbereich Mathematik und Informatik, Freie Universität Berlin.

Huang, W., Li, L., Myers, J. R., and Marth, G. T. (2012). ART: a next-generation sequencing read simulator. Bioinformatics 28, 593–594. doi: 10.1093/bioinformatics/btr708

Hume, S., Nelson, T. N., Speevak, M., McCready, E., Agatep, R., Feilotter, H., et al. (2019). CCMG practice guideline: laboratory guidelines for next-generation sequencing. J. Med. Genet. 56, 792–800. doi: 10.1136/jmedgenet-2019-106152

Ioannidis, N. M., Rothstein, J. H., Pejaver, V., Middha, S., McDonnell, S. K., Baheti, S., et al. (2016). REVEL: an ensemble method for predicting the pathogenicity of rare missense variants. Am. J. Hum. Genet. 99, 877–885. doi: 10.1016/j.ajhg.2016.08.016

Jennings, L. J., Arcila, M. E., Corless, C., Kamel-Reid, S., Lubin, I. M., Pfeifer, J., et al. (2017). Guidelines for validation of next-generation sequencing–based oncology panels: a joint consensus recommendation of the Association for Molecular Pathology and College of American Pathologists. J. Mol. Diagn. 19, 341–365. doi: 10.1016/j.jmoldx.2017.01.011

Krusche, P., Trigg, L., Boutros, P. C., Mason, C. E., De La Vega, F. M., Moore, B. L., et al. (2019). Best practices for benchmarking germline small-variant calls in human genomes. Nat. Biotechnol. 37, 555–560. doi: 10.1038/s41587-019-0054-x

Landrum, M. J., Lee, J. M., Benson, M., Brown, G. R., Chao, C., Chitipiralla, S., et al. (2018). ClinVar: improving access to variant interpretations and supporting evidence. Nucleic Acids Res. 46, D1062–D1067. doi: 10.1093/nar/gkx1153

Lavrichenko, K., Engdal, E. S., Marvig, R. L., Jemt, A., Vignes, J. M., Almusa, H., et al. (2025). Recommendations for bioinformatics in clinical practice. Genome Med. 17, 124. doi: 10.1186/s13073-025-01543-4

Li, Z., Fang, S., Zhang, R., Yu, L., Zhang, J., Bu, D., et al. (2021). VarBen: generating in silico reference data sets for clinical next-generation sequencing bioinformatics pipeline evaluation. J. Mol. Diagn. 23, 285–299. doi: 10.1016/j.jmoldx.2020.11.010

McLaren, W., Gil, L., Hunt, S. E., Riat, H. S., Ritchie, G. R. S., Thormann, A., et al. (2016). The Ensembl Variant Effect Predictor. Genome Biol. 17, 122. doi: 10.1186/s13059-016-0974-4

Mercer, T. R., Xu, J., Mason, C. E., Tong, W., and MAQC/SEQC2 Consortium (2021). The Sequencing Quality Control 2 study: establishing community standards for sequencing in precision medicine. Genome Biol. 22, 306. doi: 10.1186/s13059-021-02528-3

Mighton, C., Smith, A. C., Mayers, J., Tomaszewski, R., Taylor, S., Hume, S., et al. (2022). Data sharing to improve concordance in variant interpretation across laboratories: results from the Canadian Open Genetics Repository. J. Med. Genet. 59, 571–578. doi: 10.1136/jmedgenet-2021-107738

Milhaven, M., and Pfeifer, S. P. (2023). Performance evaluation of six popular short-read simulators. Heredity 130, 55–63. doi: 10.1038/s41437-022-00577-3

Miller, D. T., Lee, K., Gordon, A. S., Amendola, L. M., Adelman, K., Bale, S. J., et al. (2021). Recommendations for reporting of secondary findings in clinical exome and genome sequencing, 2021 update: a policy statement of the American College of Medical Genetics and Genomics (ACMG). Genet. Med. 23, 1391–1398. doi: 10.1038/s41436-021-01171-4

Miller, D. T., Lee, K., Abul-Husn, N. S., Amendola, L. M., Brothers, K., Chung, W. K., et al. (2023a). ACMG SF v3.2 list for reporting of secondary findings in clinical exome and genome sequencing: a policy statement of the American College of Medical Genetics and Genomics (ACMG). Genet. Med. 25, 100866. doi: 10.1016/j.gim.2023.100866

Miller, T. L. A., Conceição, H. B., Mercuri, R. L., Santos, F. R. C., Barreiro, R., Buzzo, J. L., et al. (2023b). Sandy: a user-friendly and versatile NGS simulator to facilitate sequencing assay design and optimization. bioRxiv [Preprint]. doi: 10.1101/2023.08.25.554791

Mitev, V. (2026). A methodological framework for real-world performance studies of clinical variant classification platforms at early organizational stages. Front. Genet. 17, 1925492. doi: 10.3389/fgene.2026.1925492

Morales, J., Pujar, S., Loveland, J. E., Astashyn, A., Bennett, R., Berry, A., et al. (2022). A joint NCBI and EMBL-EBI transcript set for clinical genomics and research. Nature 604, 310–315. doi: 10.1038/s41586-022-04558-8

Mu, J. C., Mohiyuddin, M., Li, J., Bani Asadi, N., Gerstein, M. B., Abyzov, A., et al. (2015). VarSim: a high-fidelity simulation and validation framework for high-throughput genome sequencing with cancer applications. Bioinformatics 31, 1469–1471. doi: 10.1093/bioinformatics/btu828

Patil, S. A., Mujacic, I., Ritterhouse, L. L., Segal, J. P., and Kadri, S. (2019). insiM: in silico mutator software for bioinformatics pipeline validation of clinical next-generation sequencing assays. J. Mol. Diagn. 21, 19–26. doi: 10.1016/j.jmoldx.2018.08.001

Poplin, R., Chang, P. C., Alexander, D., Schwartz, S., Colthurst, T., Ku, A., et al. (2018). A universal SNP and small-indel variant caller using deep neural networks. Nat. Biotechnol. 36, 983–987. doi: 10.1038/nbt.4235

Preston, C. G., Wright, M. W., Madhavrao, R., Harrison, S. M., Goldstein, J. L., Luo, X., et al. (2022). ClinGen Variant Curation Interface: a variant classification platform for the application of evidence criteria from ACMG/AMP guidelines. Genome Med. 14, 6. doi: 10.1186/s13073-021-01004-8

Rehder, C., Bean, L. J. H., Bick, D., Chao, E., Chung, W., Das, S., et al. (2021). Next-generation sequencing for constitutional variants in the clinical laboratory, 2021 revision: a technical standard of the American College of Medical Genetics and Genomics (ACMG). Genet. Med. 23, 1399–1415. doi: 10.1038/s41436-021-01139-4

Richards, S., Aziz, N., Bale, S., Bick, D., Das, S., Gastier-Foster, J., et al. (2015). Standards and guidelines for the interpretation of sequence variants: a joint consensus recommendation of the American College of Medical Genetics and Genomics and the Association for Molecular Pathology. Genet. Med. 17, 405–424. doi: 10.1038/gim.2015.30

Riggs, E. R., Andersen, E. F., Cherry, A. M., Kantarci, S., Kearney, H., Patel, A., et al. (2020). Technical standards for the interpretation and reporting of constitutional copy-number variants: a joint consensus recommendation of the American College of Medical Genetics and Genomics (ACMG) and the Clinical Genome Resource (ClinGen). Genet. Med. 22, 245–257. doi: 10.1038/s41436-019-0686-8

Roy, S., Coldren, C., Karunamurthy, A., Kip, N. S., Klee, E. W., Lincoln, S. E., et al. (2018). Standards and guidelines for validating next-generation sequencing bioinformatics pipelines: a joint recommendation of the Association for Molecular Pathology and the College of American Pathologists. J. Mol. Diagn. 20, 4–27. doi: 10.1016/j.jmoldx.2017.11.003

Roy, S., Tremblay, M. W., Lockhart, E., Aradhya, S., Bayrak-Toydemir, P., Bowser, M., et al. (2025). From expert knowledge to validation resources – a case for using in silico approaches to close the gap in available reference materials for common germline genetic tests. J. Mol. Diagn. 27, 1074–1083. doi: 10.1016/j.jmoldx.2025.07.006

Sola, L., Bagordo, D., Carpanzano, S., Santorsola, M., and Lescai, F. (2026). Eduomics: a Nextflow pipeline to simulate-omics data for education. NAR Genom. Bioinform. 8, lqag029. doi: 10.1093/nargab/lqag029

Stephens, Z. D., Hudson, M. E., Mainzer, L. S., Taschuk, M., Weber, M. R., and Iyer, R. K. (2016). Simulating next-generation sequencing datasets from empirical mutation and sequencing models. PLoS One 11, e0167047. doi: 10.1371/journal.pone.0167047

Stromberg, M., Roy, R., Lajugie, J., Jiang, Y., Li, H., and Margulies, E. (2017). NIRVANA: clinical grade variant annotator. In Proceedings of the 8th ACM International Conference on Bioinformatics, Computational Biology, and Health Informatics, 596. doi: 10.1145/3107411.3108204

Vasimuddin, M., Misra, S., Li, H., and Aluru, S. (2019). Efficient architecture-aware acceleration of BWA-MEM for multicore systems. In 2019 IEEE International Parallel and Distributed Processing Symposium (IPDPS), 314–324. doi: 10.1109/IPDPS.2019.00041

Wagner, J., Olson, N. D., Harris, L., Khan, Z., Farek, J., Mahmoud, M., et al. (2022a). Benchmarking challenging small variants with linked and long reads. Cell Genom. 2, 100128. doi: 10.1016/j.xgen.2022.100128

Wagner, J., Olson, N. D., Harris, L., McDaniel, J., Cheng, H., Fungtammasan, A., et al. (2022b). Curated variation benchmarks for challenging medically relevant autosomal genes. Nat. Biotechnol. 40, 672–680. doi: 10.1038/s41587-021-01158-1

Yu, Z., Du, F., Ban, R., and Zhang, Y. (2020). SimuSCoP: reliably simulate Illumina sequencing data based on position and context dependent profiles. BMC Bioinformatics 21, 331. doi: 10.1186/s12859-020-03665-5

Zhao, H., Sun, Z., Wang, J., Huang, H., Kocher, J. P., and Wang, L. (2014). CrossMap: a versatile tool for coordinate conversion between genome assemblies. Bioinformatics 30, 1006–1007. doi: 10.1093/bioinformatics/btt730

