## Supplementary for "An end-to-end in silico framework for truth-based validation and regression testing of clinical genome sequencing pipelines, from simulated reads to reportable findings"

### Supplementary Note S1. Synthetic novel variants

Question. A tertiary tool can classify a curated variant from a database assertion (PP5) rather than from the evidence. We tested whether BIAS-2015 classifies predicted loss-of-function variants that no database holds, from PVS1 and PM2.

Design. A generator script wrote the variants as an explicit list of the suite configuration, deterministically from a seed. Genes were the autosomal dominant genes of the secondary-findings list (version 3.2; Miller et al., 2023a) with a ClinGen haploinsufficiency score of 3 (ClinGen, 2026); RET and APOB were excluded because loss of function does not cause their secondary-findings condition. Of 43 eligible genes, 33 received variants. On MANE Select transcripts (release 1.5; Morales et al., 2022), the generator drew nonsense SNVs, canonical splice-site variants (±1 and ±2), and frameshift insertions and deletions of 1–20 bp (lengths not divisible by three). It kept positions where the change is predicted to trigger nonsense-mediated decay: not in the last exon, more than 50 nucleotides upstream of the last exon–exon junction, and not within the first 100 nucleotides of the coding sequence. Indels lay at least 3 bp from an exon boundary, all variants lay at least 5 bp outside homopolymer and tandem-repeat strata, and indels that would shift in a repeat were drawn again. A candidate was excluded when a ClinVar record (release of 28 September 2026), an eRepo allele (export of 15 October 2026), or a gnomAD v4.1 exome record (Chen et al., 2024) overlapped it. VEP 115 had to confirm the intended consequence on the MANE Select transcript (McLaren et al., 2016). Variants lay at least 500 bp apart, with at most 18 per gene.

A first set held 234 variants in three samples. Because ClinVar records cover almost every base of these genes, the overlap rule removed nearly all deletions, and the set contained only 5. A second set therefore planted 90 frameshift deletions (30 per length bin) in a fourth sample, excluding a deletion only when a record overlapped its breakpoints. The setting was that of Supplementary Note S2. The expected class was P or LP and the variant had to be reported. The primary score (A) requires P or LP and a report; a secondary analysis (B) also accepts a VUS with PVS1 and PM2 as the only criteria, to separate errors in the criteria from errors in combining them.

Expected class. We checked the expected class against the two current recommendations for combining criteria. The ClinGen SVI lowered PM2 to supporting strength and added the rule that one very strong and one supporting pathogenic criterion give likely pathogenic, so that novel loss-of-function variants with PVS1 and PM2 stay likely pathogenic (ClinGen Sequence Variant Interpretation Working Group, 2020). The ACGS guidelines combine criteria by points (very strong 8, strong 4, moderate 2, supporting 1; pathogenic 10 or more, likely pathogenic 6–9) but keep PM2 at moderate strength (Durkie et al., 2024). For PVS1 at very strong strength with PM2, the SVI rule gives likely pathogenic and ACGS gives 10 points, pathogenic. For PVS1 at strong strength, which BIAS-2015 assigned to 19 frameshift variants, the two disagree: the SVI rule does not cover strong plus supporting, which remains VUS under the 2015 rules, whereas ACGS gives

6 points, likely pathogenic. The expected class P or LP therefore follows both recommendations for the variants with PVS1 at very strong strength and ACGS for the others.

Results. All 324 variants were present in the true alignments and called with the correct genotype. With Nirvana, BIAS-2015 assigned PVS1 to 323 variants, at very strong strength to 304 and at strong strength to 19 frameshift variants, and PM2 at supporting strength to all 324. Only 83 variants were classified P or LP and reported (25.6%; 95% CI 21.2–30.6%; Supplementary Table S4). PVS1 with PM2 at supporting strength alone always gave VUS (226 variants; Supplementary Table S5). Every P or LP classification included a further criterion: PM1 (57 variants), PP3 at strong strength (23, all splice-site variants), or PS1 (5). Frameshift length did not change the result. Under scoring B, 309 of 324 variants were correct (95.4%; 92.5–97.2%). Of the remaining 15, 14 had also received BP4 (10 nonsense and 4 splice-site variants), and one splice-site variant in SCN5A did not receive PVS1. Switching off PP5 and BP6 did not change the class of any synthetic variant. The annotators disagreed on one deletion (chr9:99129095 CT>C; VUS with Nirvana and P with VEP).

Interpretation and limits. BIAS-2015 applied the expected criteria and the downgrade of PM2 to supporting strength recommended by the ClinGen SVI, but combined the criteria by the rules of 2015 (Richards et al., 2015) and not by the rule of the same recommendation by which PVS1 with PM2 at supporting strength is likely pathogenic (ClinGen Sequence Variant Interpretation Working Group, 2020). Under the ACGS point system, which keeps PM2 at moderate strength, variants with PVS1 at very strong strength would score 10 points, which is pathogenic, and those with PVS1 at strong strength 6 points, which is likely pathogenic (Durkie et al., 2024). Both current recommendations therefore give P or LP for nearly all of these variants, and the primary score counts the VUS as failures; the secondary analysis places the cause in the combining rule. Because the class did not change without PP5 and BP6, the result did not come from a database. Which variants reached P depended on gene-level evidence such as protein domains (PM1), not on variant type or length. The variants lie in secondary-findings genes, so the result may differ in other genes, and only one tertiary tool was tested.

### Supplementary Note S2. Indel-length experiment

Question. The reference implementation is limited to indels of 1–20 bp, and indels longer than 15 bp remain challenging to call in medically relevant genes (Wagner et al., 2022b). We tested where calling of the tested pipeline stops and whether its report decisions for real indels depend on their length.

Design. We used the indel ladder of the selection rules with the ClinVar release of 28 September 2026 and indels with at least two review stars. Length was the number of inserted or deleted bases, and the bins were 1–5, 6–10, 11–20, 21–35, 36–50, and 51–100 bp. In the autosomal dominant genes of the secondary-findings list (version 3.2; Miller et al., 2023a), P/LP indels were planted in the positive and B/LB indels in the negative sample and scored at both levels. P/LP indels in any gene filled each bin and were scored at the calling level only. The setting was that of the difficult-context test set: GIAB HG002 v5.0q background (records up to 20 bp), reads simulated at 30-fold from the GRCh38 sequence with the GIAB mask of falsely duplicated regions, two male samples, nf-core/sarek 3.10.0 with DeepVariant, and the four tertiary configurations. All 450 planted indels were present in Mason's true alignments on the intended haplotype. Calling was scored from the hap.py decision for each planted indel (true positive with the correct genotype).

Results. Every indel of 1–20 bp was called with the correct genotype (240/240), against 71 of 74 of 21–35 bp, 53 of 66 of 36–50 bp, and 12 of 70 of 51–100 bp (Supplementary Table S3). Long insertions failed earlier than long deletions: between 36 and 50 bp, 42 of 45 deletions but only 6 of 15 insertions were called. Above 50 bp, 5 of 48 deletions and 4 of 15 insertions were called. Sixteen indels, 14 of them deletion–insertions, were called in another representation; BIAS-2015 classified the parts, so no finding matched the planted indel. At the report level, 53 of 85 P/LP indels in secondary-findings genes were reported with Nirvana and with VEP, 34 with Nirvana without PP5 and BP6, and 12 with VEP without them. All 35 B/LB indels were withheld in every configuration.

Interpretation and limits. Calling of this pipeline was complete up to 20 bp and declined above 35 bp, so the scope of 1–20 bp of the reference implementation holds for it. As in the main test sets, report sensitivity for real P/LP indels depended on PP5. The bins above 20 bp contain few variants of secondary-findings genes (11–15 P/LP and 0–4 B/LB per bin), so report-level results by length are imprecise. In the first comparison, the framework's read-level check assigned 59 indels longer than 20 bp to mismapped reads although hap.py matched their calls: around a long indel the aligner clips the reads, so fewer aligned reads carry the allele than Mason's true alignments. The check now applies only to an allele that was not called, and the repeated comparison agrees with hap.py.

### Supplementary Note S3. Background-window control

Question. The test sets remove GIAB HG002 background records that overlap a planted variant or lie within 500 bp of it. Each planted variant then has an unambiguous haplotype. In somatic calling, false-positive calls were enriched near germline variants (Li et al., 2021), so the window could hide errors that a neighbouring variant would cause. We tested whether the window changes the results.

Design. We rebuilt the short1 test set (chromosomes 17 and 19; one positive and one negative male sample) with a 10 bp background window. The random seed, the planted variants, and all other settings were unchanged. Both versions were processed with the same releases of Nextflow (26.10.2), nf-core/sarek (3.10.0), references, annotation data, and container images, and with the same four tertiary configurations (BIAS-2015 with Nirvana or VEP, PP5/BP6 on or off). We then compared the outcome of every planted variant between the two windows: outcome class, failing stage, assigned class, report decision, and the call.

Results. The planted variants were identical in both versions. The 10 bp window restored 55 background records (23 in the positive and 32 in the negative sample) near 18 of the 33 planted variants; two records within 10 bp stayed removed. The nearest restored neighbour lay within 300 bp of 17 planted variants, within 150 bp, the length of one read, of 12, and within 50 bp of 2 (closest 35 bp). Every planted variant had the same outcome under both windows in all four configurations, including the 12 with a neighbour within one read length (Supplementary Table S2).

The negative sample of the control had two additional false reports. Both lay far from any planted variant and from any restored background record, and the truth in both regions was identical in the two versions. One was a true homozygous HG002 variant in LILRB3 (chr19:54220736 A>C), called at a depth of 3 in the LILR/KIR region only in the control and classified as pathogenic in all four configurations. The other was a false-positive call in PLD2 (chr17:4808374 T>A; 2 of 15 reads with the alternative allele), reported only in the VEP-based configurations. A likely cause is that the added background changed the simulated haplotypes, so Mason sampled different reads along the whole chromosome even with the same seed. We did not verify this directly.

Interpretation and limits. The window did not change the outcome of any planted variant. The control is small and covers two chromosomes, so it excludes a large effect of the window in this design but not a small one. It also does not test two variants in the same codon.

#### Supplementary Table S1. Outcomes of the difficult-context test set (difficult_v5q) by selection rule and pipeline configuration.

| Selection rule | Level | n | Nirvana | Nirvana, PP5/BP6 off | VEP | VEP, PP5/BP6 off |
|---|---|---|---|---|---|---|
| ClinVar, segmental duplications | calling | 60 | correct 60 | correct 60 | correct 60 | correct 60 |
| ClinVar, low mappability | calling | 60 | correct 60 | correct 60 | correct 60 | correct 60 |
| ClinVar, homopolymers | calling | 60 | correct 60 | correct 60 | correct 60 | correct 60 |
| ClinVar, tandem repeats | calling | 60 | correct 60 | correct 60 | correct 60 | correct 60 |
| ClinVar, GRCh38 false duplications | calling | 20 | correct 5; filtered 1; mismapped 9; not_called 5 | correct 5; filtered 1; mismapped 9; not_called 5 | correct 5; filtered 1; mismapped 9; not_called 5 | correct 5; filtered 1; mismapped 9; not_called 5 |
| ClinVar, paralogous genes (SMN1, PMS2, OTOA) | calling | 7 | correct 3; mismapped 3; not_called 1 | correct 3; mismapped 3; not_called 1 | correct 3; mismapped 3; not_called 1 | correct 3; mismapped 3; not_called 1 |
| eRepo P/LP, SF genes, difficult regions | both | 15 | correct 14; misclassified 1 | correct 12; misclassified 3 | correct 14; misclassified 1 | correct 11; misclassified 4 |
| eRepo B/LB, SF genes, difficult regions | both | 10 | correct 7; misclassified 3 | correct 5; misclassified 5 | correct 9; misclassified 1 | correct 4; misclassified 6 |

Background: GIAB HG002 v5.0q; reads simulated from GRCh38 with the GIAB mask of falsely duplicated regions; the pipeline used the standard GRCh38 reference. Outcome classes as in Table 3; calling-level variants are not classified, so their outcomes are identical in all configurations. Calling with the correct genotype: 273/292 (93.5%; 95% Wilson interval 90.1–95.8%). Report level (both-level variants): P/LP reported 14/15 (Nirvana), 12/15 (Nirvana, PP5/BP6 off), 14/15 (VEP), 11/15 (VEP, PP5/BP6 off); B/LB withheld 10/10 in all configurations. SF, ACMG secondary-findings genes, version 3.2.

#### Supplementary Table S2. Outcome of each planted variant with a 500 bp and a 10 bp background window (test set short1).

| Sample | Planted variant (GRCh38) | Gene | Nearest restored record (bp) | Nirvana | Nirvana, PP5/BP6 off | VEP | VEP, PP5/BP6 off |
|---|---|---|---|---|---|---|---|
| positive | chr17:43057127:A:C | BRCA1 | – | correct | correct | correct | correct |
| positive | chr17:43063951:T:A | BRCA1 | – | correct | misclassified | correct | misclassified |
| positive | chr17:43124027:ACT:A | BRCA1 | – | correct | correct | correct | misclassified |
| positive | chr17:7670669:G:T | TP53 | – | correct | correct | correct | correct |
| positive | chr17:7674252:C:T | TP53 | – | correct | correct | correct | correct |
| positive | chr17:7675989:C:T | TP53 | 164 | correct | correct | correct | correct |
| positive | chr17:80110024:CCAA:C | GAA | 96 | correct | misclassified | correct | misclassified |
| positive | chr17:80117059:G:AT | | 69 | not_annotated | not_annotated | not_annotated | not_annotated |
| positive | chr19:11089550:T:C | LDLR | – | correct | correct | correct | correct |
| positive | chr19:11100349:A:T | LDLR | 112 | correct | correct | correct | correct |
| positive | chr19:11102741:G:A | LDLR | – | correct | correct | correct | correct |
| positive | chr19:11105258:G:T | LDLR | – | correct | correct | correct | correct |
| positive | chr19:11107472:A:G | LDLR | – | correct | misclassified | correct | correct |

| | | | | | | | |
|---|---|---|---|---|---|---|---|
| positive | chr19:11113743:G:C | LDLR | 71 | correct | correct | correct | misclassified |
| positive | chr19:38455359:A:G | RYR1 | 147 | correct | correct | correct | correct |
| positive | chr19:38494426:G:C | RYR1 | – | correct | misclassified | correct | correct |
| positive | chr19:38499223:G:A | RYR1 | 472 | correct | correct | correct | correct |
| negative | chr17:7669671:C:G | TP53 | – | correct | misclassified | correct | misclassified |
| negative | chr17:7676567:C:G | TP53 | 83 | correct | misclassified | correct | misclassified |
| negative | chr17:80108741:C:G | GAA | 35 | correct | correct | correct | correct |
| negative | chr17:80111997:AC:A | GAA | 198 | report_mismatch | misclassified | report_mismatch | report_mismatch |
| negative | chr17:80118264:G:A | GAA | – | correct | correct | correct | correct |
| negative | chr19:11116873:C:T | LDLR | 52 | misclassified | misclassified | correct | correct |
| negative | chr19:11120432:G:C | LDLR | 226 | misclassified | misclassified | misclassified | misclassified |
| negative | chr19:11123210:C:T | LDLR | 54 | correct | correct | correct | correct |
| negative | chr19:11128126:C:A | LDLR | 253 | correct | correct | correct | correct |
| negative | chr19:38443790:G:A | RYR1 | 113 | correct | correct | correct | correct |
| negative | chr19:38473635:A:G | RYR1 | – | correct | correct | correct | correct |
| negative | chr19:38477816:A:G | RYR1 | – | correct | correct | correct | correct |
| negative | chr19:38500000:G:A | RYR1 | 35 | correct | correct | misclassified | misclassified |
| negative | chr19:38543551:A:G | RYR1 | 130 | correct | correct | misclassified | misclassified |
| negative | chr19:38561236:C:A | RYR1 | 220 | misclassified | correct | correct | misclassified |
| negative | chr19:38566986:G:C | RYR1 | – | correct | correct | correct | correct |

Outcome classes as in Table 3; each outcome was identical with the 500 bp and the 10 bp window. Nearest restored record: distance to the closest background record that the 10 bp window kept and the 500 bp window removed; –, none. Additional false reports in the control (not planted): LILRB3 chr19:54220736 A>C (all four configurations) and PLD2 chr17:4808374 T>A (VEP configurations).

### Supplementary Table S3. Calling and report decisions by indel length (indel-length experiment).

| Length (bp) | Planted | Called, correct genotype (95% CI) | Deletions | Insertions | Deletion–insertions | P/LP reported: Nirvana | Nirvana, PP5/BP6 off | VEP | VEP, PP5/BP6 off | B/LB withheld (all) |
|---|---|---|---|---|---|---|---|---|---|---|
| 1–5 | 80 | 80/80 = 100.0% (95.4–100.0) | 58/58 | 20/20 | 2/2 | 13/15 | 9/15 | 13/15 | 3/15 | 10/10 |
| 6–10 | 80 | 80/80 = 100.0% (95.4–100.0) | 53/53 | 21/21 | 6/6 | 11/15 | 6/15 | 11/15 | 5/15 | 10/10 |
| 11–20 | 80 | 80/80 = 100.0% (95.4–100.0) | 56/56 | 11/11 | 13/13 | 10/15 | 5/15 | 10/15 | 1/15 | 10/10 |
| 21–35 | 74 | 71/74 = 95.9% (88.7–98.6) | 51/51 | 13/15 | 7/8 | 11/15 | 6/15 | 11/15 | 1/15 | 4/4 |
| 36–50 | 66 | 53/66 = 80.3% (69.2–88.1) | 42/45 | 6/15 | 5/6 | 7/11 | 7/11 | 7/11 | 2/11 | – |
| 51–100 | 70 | 12/70 = 17.1% (10.1–27.6) | 5/48 | 4/15 | 3/7 | 1/14 | 1/14 | 1/14 | 0/14 | 1/1 |
| All | 450 | 376/450 = 83.6% (79.8–86.7) | 265/311 | 75/97 | 36/42 | 53/85 | 34/85 | 53/85 | 12/85 | 35/35 |

Length: inserted or deleted bases (longer allele minus the shared anchor base for deletion–insertions). Called: hap.py true positive with the correct genotype; 95% Wilson score intervals. P/LP reported and B/LB withheld: variants in secondary-findings genes scored at both levels (P/LP 15, 15, 15, 15, 11, and 14 per bin; B/LB 10, 10, 10, 4, 0, and 1); B/LB results were identical in the four configurations; –, no variant.

### Supplementary Table S4. Synthetic variants by type, scored with two expected classes (configuration Nirvana).

| Variant type | n | Primary (A): P or LP, reported (95% CI) | Secondary (B): A, or VUS from PVS1 and PM2 alone (95% CI) |
|---|---|---|---|
| Nonsense | 30 | 10/30 = 33.3% (19.2–51.2) | 20/30 = 66.7% (48.8–80.8) |
| Canonical splice site | 30 | 23/30 = 76.7% (59.1–88.2) | 25/30 = 83.3% (66.4–92.7) |
| Frameshift insertion, 1–5 bp | 54 | 9/54 = 16.7% (9.0–28.7) | 54/54 = 100.0% (93.4–100.0) |
| Frameshift insertion, 6–10 bp | 56 | 10/56 = 17.9% (10.0–29.8) | 56/56 = 100.0% (93.6–100.0) |
| Frameshift insertion, 11–20 bp | 59 | 15/59 = 25.4% (16.1–37.8) | 59/59 = 100.0% (93.9–100.0) |
| Frameshift deletion, 1–5 bp | 35 | 4/35 = 11.4% (4.5–26.0) | 35/35 = 100.0% (90.1–100.0) |
| Frameshift deletion, 6–10 bp | 30 | 7/30 = 23.3% (11.8–40.9) | 30/30 = 100.0% (88.6–100.0) |
| Frameshift deletion, 11–20 bp | 30 | 5/30 = 16.7% (7.3–33.6) | 30/30 = 100.0% (88.6–100.0) |
| All | 324 | 83/324 = 25.6% (21.2–30.6) | 309/324 = 95.4% (92.5–97.2) |

Scoring A requires the expected class (P or LP) and a report; scoring B also accepts a VUS whose only criteria were PVS1 and PM2. The four configurations gave the same classes except for one deletion (see text). 95% Wilson score intervals.

### Supplementary Table S5. ACMG/AMP criteria assigned by BIAS-2015 to the synthetic variants (configuration Nirvana).

| **Criteria assigned** | **Class** | **Nonsense** | **Splice site** | **Frameshift insertion** | **Frameshift deletion** | **All** |
|---|---|---|---|---|---|---|
| PM2_Supporting + PVS1 | VUS | 10 | 2 | 121 | 76 | 209 |
| PM1_Moderate + PM2_Supporting + PVS1 | P | 4 | – | 22 | 11 | 37 |
| PM2_Supporting + PP3_Strong + PVS1 | P | – | 19 | – | – | 19 |
| PM2_Supporting + PVS1_Strong | VUS | – | – | 14 | 3 | 17 |
| BP4_Supporting + PM2_Supporting + PVS1 | VUS | 9 | 4 | – | – | 13 |
| PM1_Strong + PM2_Supporting + PVS1 | P | – | – | 6 | 3 | 9 |
| PM1_Supporting + PM2_Supporting + PVS1 | P | – | – | 2 | 2 | 4 |
| BP4_Supporting + PM1_Moderate + PM2_Supporting + PVS1 | P | 3 | – | – | – | 3 |
| BP4_Supporting + PM2_Supporting + PP3_Strong + PVS1 | P | – | 3 | – | – | 3 |
| PM2_Supporting + PS1_Moderate + PVS1 | P | 2 | – | 1 | – | 3 |
| BP4_Strong + PM2_Supporting + PVS1 | VUS | 1 | – | – | – | 1 |
| BP4_Supporting + PM1_Strong + PM2_Supporting + PVS1 | P | 1 | – | – | – | 1 |
| PM1_Moderate + PM2_Supporting + PP3_Strong + PVS1 | P | – | 1 | – | – | 1 |
| PM1_Moderate + PM2_Supporting + PVS1_Strong | LP | – | – | 1 | – | 1 |
| PM1_Strong + PM2_Supporting + PS1_Moderate + PVS1 | P | – | – | 1 | – | 1 |
| PM2_Supporting | VUS | – | 1 | – | – | 1 |
| PM2_Supporting + PS1_Moderate + PVS1_Strong | LP | – | – | 1 | – | 1 |

Criteria as reported by BIAS-2015 with their applied strength; PVS1 without suffix is very strong. –, none.

### References

All other works cited in this Supplementary Material are listed in the References of the main article.

ClinGen (2026). ClinGen dosage sensitivity curation: gene curation list, GRCh38. Available at: https://ftp.clinicalgenome.org/ClinGen_gene_curation_list_GRCh38.tsv (accessed 28 October 2026).
